# Key transcription factors influence the epigenetic landscape to regulate retinal cell differentiation

**DOI:** 10.1101/2022.03.05.483140

**Authors:** Yichen Ge, Xushen Chen, Nan Nan, Jonathan Bard, Fuguo Wu, Donald Yergeau, Tao Liu, Jie Wang, Xiuqian Mu

**Author notes:** **To whom correspondence should be addressed:** Xiuqian Mu,; Jie Wang,.

## Abstract

How the diverse neural cell types emerge from multipotent neural progenitor cells during central nervous system development remains poorly understood. Recent scRNA-seq studies have delineated the developmental trajectories of individual neural cell types in many neural systems including the neural retina. Further understanding of the formation of neural cell diversity requires knowledge about how the epigenetic landscape shifts along individual cell lineages and how key transcription factors regulate these changes. In this study, we dissect the changes in the epigenetic landscape during early retinal cell differentiation by scATAC-seq and identify globally the enhancers, enriched motifs, and potential interacting transcription factors underlying the cell state/type specific gene expression in individual lineages. Using CUT&Tag, we further identify the enhancers bound directly by four key transcription factors, Otx2, Atoh7, Pou4f2, and Isl1, and uncover their roles in shaping the epigenetic landscape and controlling gene expression in a sequential and combinatorial fashion along individual retinal cell lineages such as retinal ganglion cells (RGCs). Our results reveal a general paradigm in which transcription factors collaborate and compete to regulate the emergence of distinct retinal cell types such as RGCs from multipotent retinal progenitor cells (RPCs).

## INTRODUCTION

How diverse neural cell types emerge from multipotent neural progenitor cells during central nervous system development remains poorly understood. We address this question in the neural retina. The retina functions to receive visual signals, transform them into electrophysiological pulses, and transmit them to the brain. This task is carried out by the various retinal neurons, which form the visual circuitry (1, 2). All retinal cell types, including photoreceptors (rods and cones), retinal ganglion cells (RGCs), horizontal cells, amacrine cells, bipolar cells, and Müller glial cells, are generated from multipotent retinal progenitor cells (RPCs) during development (3). Retinal cell differentiation involves several aspects (3–5). First, the different retinal cell types are generated in a conserved temporal order with RGCs always the first cell type to form, and Müller cells the last. On the other hand, there are significant temporal overlaps in the genesis of the different retinal cell types, and several retinal cell types often form concurrently (6). Two major waves of cell differentiation occur during retinal development, the first including RGCs, horizontal cells, amacrine cells, and cones, and the second including rods, bipolar cells, and Müller cells. Therefore, at almost any given time during retinal development, RPCs have to adopt one of the several cell fates. The prevailing model for the temporally sequential genesis of retinal cell types is that RPCs undergo a series of competence changes, so that RPCs at early developmental stages only generate the early cell types, whereas late RPCs only generate the late cell types (4, 7). Although some factors influencing RPC competence have been identified (8–12), the genetic and molecular nature underlying RPC competence has not been definitively defined. Nevertheless, each retinal lineage follows a distinct developmental trajectory in the stepwise process from naïve RPCs to differentiated cells and is controlled by a distinct genetic program; regulatory cascades composed of multiple regulators for individual retinal cell types have been identified and continue to be elaborated (13–17).

Much evidence suggests that RPCs are heterogeneous (18, 19), but recent single-cell transcriptomics studies indicate that the different cell lineages all go through a shared state of transitional RPCs (tRPCs) following individual developmental trajectories (8, 20–23). These studies have provided an unprecedented perspective on the cellular process of retinal cell differentiation by clearly and unequivocally delineating the phases of the individual trajectories and associated genes. For example, the RGC trajectory starts from naïve RPCs (nRPCs), goes through tRPCs, and then reaches fate committed (early) and eventually differentiated (late) RGCs. Importantly, tRPCs co-express transcription factor genes for different cell fates, such as *Atoh7* and *Neurog2* for RGCs, *Foxn4* for horizontal cells and amacrine cells (H&As), and *Otx2* and *Neurod1* for photoreceptors (PHCs) (21, 22). Further, tRPCs display downregulated Notch signaling and upregulated expression of Notch ligands, remain multipotent, are ready to exit or have just exited the cell cycle, and eventually assume one of the retinal cell fates. These findings suggest that tRPCs are the competent RPCs and that their competence for the different cell types is determined by the transcription factors co-expressed in them at a given developing time (22). It has been proposed that transcription factors for different lineages compete to drive tRPCs to distinct fates, but the mechanisms by which such competition takes place are not known (22).

RGCs are the only output neurons in the retina, which synapse with upstream neurons in the visual circuitry and project axons to the brain via the optic nerve (24, 25). Although there are more than 40 types of RGCs in the mouse as determined by anatomy, physiology, ad single cell transcriptomics (26–28), their genesis is subject to the regulation of a shared genetic pathway, as the mutation of a single gene *Atoh7* leads to almost complete RGC loss (29, 30). RGCs are generated in the first wave of retinal cell differentiation, overlapping with cone, horizontal cell, and amacrine cell genesis; in the mouse, the formation of RGCs starts at around E11, reaches the peak at E14.5, and completes at around birth (6). Like other retinal cell types, RGC formation is subject to a gene regulatory cascade with key transcription factors functioning at different stages (14). Among these transcription factors, Atoh7 participates in the initial activation of the RGC expression program but is not solely required for establishing the RGC lineage, since the RGC lineage still arises without Atoh7, although these cells die out immediately (22, 31). Based on expression patterns and knockout phenotypes, the SoxC transcription factors, including Sox4, Sox11, and Sox12, likely function together with Atoh7 in shepherding tRPCs to the RGC fate, but this remains to be definitively established (22, 32, 33). Downstream are Pou4f2 and Isl1, two transcription factors expressed coincidentally with the RGC fate specification. Pou4f2 and Isl1, likely in collaboration with additional factors, specify the RGC fate by activating and stabilizing the RGC-specific expression program (34). Pou4f2 and Isl1, together with a large set of downstream transcription factors, are also involved in RGC maturation and type formation by regulating genes required for general RGC structure and function as well as RGC type specific features (35–41).

Conceivably, transcription factors define individual cell states/types in the different trajectories and promote the progression of different lineages by interacting with and thereby influencing the epigenetic landscape. The epigenetic landscape can be surveyed by examining the distribution of histone markers for specific chromatin states with ChIP-seq, the degrees of chromatin openness by accessibility assays such as Dnase-seq, FAIRE-seq, and ATAC-seq, or expression of enhancer RNAs (42–45). Using these techniques, several studies have been conducted to systematically analyze the epigenetic statuses at the whole retinal level across different developmental stages (46–50). Whereas the results from these studies reveal the temporal epigenetic shifts and associated gene expression changes throughout development, they lack the necessary cellular resolution to decipher how epigenetic landscape shifts along individual lineage trajectories. To address this issue, we used scATAC-seq (51) to survey the dynamic epigenetic landscape in both wild-type and *Atoh7*-null retinas at two early developmental stages (E14.5 and E17.5). Our study reveals globally the enhancers, enriched DNA motifs, and possible interacting transcription factors underlying the progression of cell state/type-specific gene expression along the trajectories of individual lineages, including that of the RGC lineage. Further, we performed CUT&Tag to identify the binding sites, and thereby the associated enhancers and the target genes, of four key transcription factors, Otx2, Atoh7, Pou4f2, and Isl1, in the E14.5 retina. Analysis of the associated enhancers and target genes of Atoh7 and Otx2 supports the idea that transcription factors for different lineages compete and/or collaborate to drive tRPCs toward distinct retinal cell fates. The enhancers bound by Atoh7, Pou4f2, and Isl1 and the associated target genes allow us to uncover how these key transcription factors regulate the epigenetic landscape in a sequential and combinatorial fashion to control gene expression during RGC genesis. Overall, these results provide a comprehensive view of the shifting epigenetic landscape along the developmental trajectories, particularly that of RGCs, in early retinal development, and reveal the roles of key transcription factors in interacting and influencing the epigenetic landscape to promote RGC genesis.

## MATERIAL AND METHODS

### Animals

The *Atoh7*^lacZ^, *Atoh7*^*zsGreenCreERT2*^ (referred to as *Atoh7*^*zsGreen*^ in the text), *Pou4f2*^*FLAGtdTomato*^ (referred to as *Pou4f2*^*tdTomato*^ in the text), *Atoh7*^*HA*^, and *Pou4f2*^*HA*^ alleles have been described previously and were maintained in the C57B/L6x 129 genetic background (30, 52, 53). *Atoh7*^*lacZ*^ and *Atoh7*^*zsGreen*^ are null alleles. *Pou4f2*^*tdTomato*^ is a wild-type allele expressing a FLAG-tagged version of Pou4f2 and tdTomato. *Atoh7*^*HA*^ and *Pou4f2*^*HA*^ alleles are phenotypically wild type. The *Pou4f2* enhancer deletion alleles were generated by CRISPR in the C57BL/6 background as described in detail below. All animal experiments were approved by the Institutional Animal Care and Use Committees of Roswell Park Comprehensive Cancer Center and the University at Buffalo. All procedures were conformed to the US Public Health Service Policy on Humane Care and Use of Laboratory Animals.

### Tissue collection, dissociation, and cell sorting

Retinas with the desired genotypes at different developmental stages (E14.5, and E17.5) were collected in cold phosphate buffered solution (PBS) after timed mating. They were then dissociated into single cell suspensions following the procedure described before (22, 53). Briefly, retinas were incubated in PBS with 100 μg/mL trypsin for 5 minutes at 37ºC and then triturated five times with a pipette. The soybean trypsin inhibitor was added to a final concentration of 100 μg/mL to stop digestion. Cells were collected by centrifugation at 500 g for 5 minutes, washed twice with PBS, and re-suspended in PBS. Cell sorting was performed on a BD FACS Fusion Cell Sorter as previously described (53). The viable and single cell events were selected for sorting through the forward and side scatter gates. The zsGreen signal was collected by a 488 laser with a 530/30 detector, and a 561 laser with a 582/15 detector was used to collect tdTomato expressing cells. The gating threshold for the fluorescence was set to relatively low levels to isolate cells that had just begun to express the two fluorescent proteins and to obtain overlapping cell populations (**Suppl. Figure 1**). The viability of purified cells was tested by trypan blue staining, and each sample had at least 80% of cells viable.

### scRNA-seq preparation and sequencing

scRNA-seq was performed on 10X Chromium single cell platform using the Chromium Single Cell 3’ Library & Gel Bead Kit v3.1, in keeping with the manufacturer’s instructions (10X Genomics, CG000204). The sorted cells were loaded onto the 10X Genomics Chromium Controller targeting >5,000 cells per sample to generate scRNA-seq libraries. The quantity and quality of the libraries were evaluated by the Qubit Fluorometer and Agilent Fragment Analyzer. High-quality libraries were subsequently sequenced on a NovaSeq6000 sequencer (PE 26×100).

### Nuclei isolation, scATAC-seq library preparation, and sequencing

Nuclei isolation was performed following the protocol from 10X Genomics (CG000169). Briefly, 100,000-1,000,000 cells were collected by FACS and centrifuged at 300 g for 5 min at 4°C. Cell pellets were resuspended in 100 μl cold Lysis Buffer (10 mM Tris-HCl pH 7.4, 10 mM NaCl, 3 mM MgCl_2_, 0.1% Tween-20, 0.1% Nonidet P40, 0.01% Digitonin, 1%BSA), mixed by gently by pipetting then incubated for 2 min on ice. One milliliter of cold Wash Buffer (10 mM Tris-HCl pH 7.4, 10 mM NaCl, 3 mM MgCl_2_, 0.1% Tween-20, 1%BSA) was then added immediately and mixed by gentle pipetting. The cells were then spun down at 500 g for 5 min at 4°C, and resuspended in cold 1X Nuclei Buffer (10x Genomics, PN-2000153). The nuclei concentration and quality were determined with a hemocytometer.

scATAC-seq library preparation was performed following the protocol from 10X Genomics (CG000168 Rev B) using the Chromium Single Cell ATAC Reagent Kits (10X Genomics). The libraries were cleaned using SPRIselect reagent (Beckman Coulter). The quantity and quality of the libraries were evaluated by the Qubit Fluorometer and Agilent Fragment Analyzer. The libraries were subsequently sequenced on a NovaSeq6000 (PE50).

### scATAC-seq data analysis

#### Data processing and clustering

Analysis of the scATAC-seq data was performed using the ArchR package (54). Raw sequence data were first processed by CellRanger ATAC (version 1.2.0) for read filtering, alignment against mouse genome (mm10), and barcode counting. The barcoded and aligned fragments results were then used as input data for analysis by ArchR (version 1.0.1), and the different samples were pooled according to development stages (E14.5 and E17.5) or all together. QC analysis was then performed to filter out low quality cells with transcription start site (TSS) enrichment score less than 4 and the number of fragments less than 1,000, and to remove doublets. A genome-wide tile matrix with insertion counts was calculated on 500-bp non-overlapping windows. Dimensional reduction was implemented by Iterative Latent Semantic Indexing (LSI) method with default two iterations, the clusters were then identified by a shared nearest neighbor (SNN) modularity optimization based clustering algorithm. Uniform Manifold Approximation and Projection (UMAP) was then performed for visualization in reduced dimension space.

#### Gene activity analysis

The GeneScore matrix was generated using distance-weighted accessibility models with the 500bp-bin tile matrix and gene window (100 kb on either side of the gene). To identify the marker genes, gene expression in each cell type was compared with bias-matched background cells using Wilcoxon test with log2 fold change higher than 1.25 and false discovery rate (FDR) less than 0.01. Genes with different gene scores between two clusters were similarly identified. We then imputed gene scores by MAGIC (55) for visualization.

#### Identification of differentially accessible peaks

Insertion counts from individual cells in each cluster were aggregated, which were considered pseudo-bulk replicates. Peak calling was performed on the pseudo-bulk replicates using MACS2 (56), and an iterative overlap peak merging procedure was then applied on the 501-bp fixed-width peaks to generate a reproducible peak set. Marker peaks were identified by comparing each cluster cells with bias-match background cells using Wilcoxon test with log2 fold change higher than 0.5 and FDR less than 0.1. To identify the differentially accessible peaks between wild-type and *Atoh7*-null cells, we focused on the tRPC clusters and the early RGC clusters and used the method described above.

#### Integrative analysis (peak to gene linkage analysis, P2G) of scATAC-seq and scRNA-seq

scRNA-seq data analysis from FACS enriched E14.5 and E17.5 retinal cells were performed as previously described (22, 53). Unconstrained integration was applied to the scATAC-seq with matched scRNA-seq by comparing the scATAC-seq gene score with the scRNA-seq gene expression matrix. A GeneIntegrationMatrix was created with pseudo-scRNA-seq profile for each scATAC-seq cell. With the peak matrix and GeneIntegrationMatrix, we computed the Pearson correlation of all possible peak-to-gene combinations within 250 kb. Only the links with correlations higher than 0.45 were visualized.

#### Trajectory analysis

We performed supervised trajectory analysis by ArchR. In our study, we focused on the RGC trajectory which was defined based on the identities of the cell clusters relevant to this lineage, including nRPCs, tRPCs, early RGCs, and late RGCs, which provided a trajectory backbone. An RGC trajectory was created based on the average positions of each cell cluster within n-dimension subspace, and individual cells were then aligned to this trajectory by calculating the nearest cell-to-trajectory distance. To identify the top variable genes or peaks along the trajectory, ArchR smoothed the gene or peak matrix using smoothing window, and the heatmaps with genes or peaks were plotted with variance quantile cutoff higher than 0.9.

#### Transcription factors and motif analysis

We used the chromVAR pipeline (57) to obtain deviation scores for motif (CIS-BP motif database) enrichment on a per-cell basis. ATAC-seq allows for the unbiased identification of transcription factors that exhibit large changes in chromatin accessibility at sites containing their DNA binding motifs. To pin down the relevant transcription factors, we generated a list of 47 transcription factors expressed at substantial levels in the developing retina based on previously published RNA-seq data (22), and only motifs recognized by these transcription factors were displayed.

#### Footprinting analysis

Cells were grouped by clusters to get pseudo-bulk ATAC-seq profiles in order to accurately profile transcription footprints. The footprints were then computed for selected motifs. To account for the Tn5 insertion bias, a k-mer frequency matrix that contains all possible k-mers across -/+ 250 bp at the motif center was created. The normalized footprint profiles were plotted by subtracting the Tn5 bias within -/+ 250bp around the motif sites.

### CUT&Tag experiment and data analysis

The Cleavage Under Targets & Tagmentation (CUT&Tag) experiments (58) were performed using reagents and following a protocol from EpiCypher. For Atoh7 and Pou4f2, anti-HA (rabbit, 3724, Cell Signaling) was used with retinal tissues from E14.5 *Atoh7*^*HA/HA*^ and *Pou4f2* ^*HA/HA*^ embryos (Fu, X., et al. 2009). The other antibodies included Anti-Otx2 (rabbit, HPA000633, Sigma) and Anti-Isl1 (rabbit, AB4326, Millipore). Control normal rabbit IgG was from Santa Cruz (SC-2027). E14.5 retinas were harvested, dissociated, and cells were collected as described above. For each experiment, nuclei were isolated from 1×10^5^ cells by incubating them in the NE Buffer (20 mM HEPES-KOH, pH 7.9, 10 mM KCl, 0.1% Triton X-100, 20% Glycerol, 0.5 mM Spermidine, 1x Roche cOmpleteTM Protease Inhibitor) for 10 min on ice. Concanavalin A-coated magnetic beads (10 μl/sample, Bangs Laboratories, #BP531) were activated in cold bead activation buffer (20 mM HEPES, pH 7.9, 10 mM KCl, 1 mM CaCl_2_,1 mM MnCl_2_). The nuclei were bound to the activated beads by incubating at room temperature for 10 min. The nuclei-beads were then resuspended in 50 μl cold Antibody Buffer (20 mM HEPES, pH 7.5, 150 mM NaCl, 0.5 mM Spermidine, 1x Protease Inhibitor, 0.01% Digitonin, 2 mM EDTA). 0.5 μg primary antibody or control IgG was then added to each sample and incubated on a nutator at 4 ºC overnight. The beads were then isolated by a magnet, the Antibody Buffer was removed, and the beads were resuspended in 50 μl cold Digitonin Buffer (20 mM HEPES, pH 7.5, 150 mM NaCl, 0.5 mM Spermidine, 1x Protease Inhibitor, 0.01% Digitonin). Then 0.5 μg anti-rabbit IgG secondary antibody (EpiCypher, 13-0047) was added to each sample and incubated on a nutator for 30 min at room temperature. Samples were washed two times in cold Digitonin Buffer and resuspended in 50 μl cold Digitonin 300 Buffer (20 mM HEPES, pH 7.5, 300 mM NaCl, 0.5 mM Spermidine, 1x Protease Inhibitor, 0.01% Digitonin). Then 2.5 μl CUTANA pAG-Tn5 (EpiCypher, #15-1017) was added and incubated on a nutator for 1 hour at room temperature. Samples were then washed twice in cold Digitonin 300 Buffer and resuspended in 50 μl cold Tagmentation Buffer (20 mM HEPES, pH 7.5, 300 mM NaCl, 0.5 mM Spermidine, 1x Protease Inhibitor, 0.01% Digitonin, 10 mM MgCl_2_), and incubated on a nutator for 1 hour at 37 ºC. After incubation, the supernatant was removed and the sample beads were resuspended in 50 μl TAPS Buffer (10 mM TAPS, pH 8.5, 0.2 mM EDTA) to stop the tagmentation reaction. After removing TAPS Buffer, 5 μl SDS Release Buffer (10 mM TAPS, pH 8.5, 0.1% SDS) was added and incubated at 58 ºC for 1 hour. Fifteen microliters of SDS Quench Buffer (0.67% Triton-X 100) was then added. CUT&Tag libraries were then generated by PCR amplification. Two microliters of uniquely barcoded i5 and i7 primers (10 μM) and 25 μl CUTANA High Fidelity 2x PCR Master Mix (EpiCypher, #15-1018) were added to each sample and mixed. PCR was performed using the following conditions: 72 ºC for 5 min, 98 ºC for 30 sec, and then twenty cycles of 98 ºC for 10 sec, 63 ºC for 10 sec, followed by an extra 1 min extension at 72 ºC. The PCR product was cleaned up using AMPure XP beads (Beckman Coulter, #A63880) following the manufacturer’s instructions. DNA was eluted into 15 μl water and quantified using the Qubit fluorometer (Invitrogen, v3.0). Library fragments were analyzed on an Agilent Fragment Analyzer and then sequenced on an Illumina NovaSeq 6000 sequencer (PE50).

To analyze the CUT&Tag data, sequence reads were aligned to the mm10 mouse reference genome using bwa-mem (59). Peaks were called under pair-end mode, with minimum FDR cutoff threshold set to 0.01, except for Isl1 which was set to 0.2 to increase the number of peaks, with MACS2 Peakcall version MACS/2.2.7.1 (56). Two independent replicates for each transcription factor were first analyzed by MACS2 and correlation of the two replicates was calculated using ucsc-bigwigCorrelate tool restricted to bigBed of merged samples to ensure reproducibility, but the eventual peak calling was performed with merged sequence reads from both replicates.

Intersection analysis of scATAC-seq peaks and CUT&Tag peaks and the resulting heatmaps were created with Deeptools/2.3.6 (60). Heatmaps were based on normalized values of E14.5 scATAC-seq peaks which overlapped with the CUT&Tag peaks for individual transcriptions factors, and gene annotation with these peaks was based on the P2G link analysis. In addition, intersecting peaks between different transcription factors were also identified based on their intersections with the E14.5 scATAC-seq data, and heatmaps were similarly generated based on scATAC-seq values using Deeptools/2.3.6 (see Results section for details). Tables associated with the CUT&Tag peaks that intersect with E14.5 scATAC-seq peaks were created utilizing bedtools/2.23.0.

### Deletion of Pou4f2 enhancers by CRISPR/Cas9, Immunofluorescence Staining and RNAscope in situ Hybridization

Two or four crRNAs targeting the flanking sequences of the enhancer region to be deleted were designed using CRISPOR (61) in combination with IDT’s web design tool, and were synthesized by IDT. The crRNAs, together with the tracrRNA and Cas9 protein were premixed to form RNPs and injected into fertilized C57BL/6 oocytes, which were then transferred to the uterus of peudopregnant female mice. After the pups were born, those carrying the desired deletions were identified by PCR using tail DNA and primers specific for deleted and wild-type alleles. Mouse lines carrying the deleted alleles were further established by breeding with wild-type C57BL/6 mice. The exact junction sequences were obtained by Sanger sequencing of the PCR product. Four enhancer deletion lines were generated including *Pou4f2*^*ER1*^, *Pou4f2*^*ER2*^, *Pou4f2*^*ER3*^, and *Pou4f2*^*ER2/3*^. The target sequences of the crRNAs, the genotyping primers, and the genomic coordinates of the deleted regions can be found in the **Suppl. Materials**.

In situ hybridization was performed using RNAscope double Z probes (Advanced Cell Diagnostics) on paraffin-embedded retinal sections. After timed mating, embryos of desired stages were collected, fixed with 4% paraformaldehyde, embedded in paraffin, sectioned at 6 μm, and de-waxed with methanol (34, 37, 39, 62). The sections were then processed, hybridization was performed, and the signals were visualized using the RNAscope® 2.5 HD Detection Reagents-RED following the manufacturer’s manual. In situ images were collected using a Nikon 80i Fluorescence Microscope equipped with a digital camera. If necessary, the contrast of images was adjusted by Adobe Photoshop to the same degree to all images in the same experiment.

## RESULTS

### Experimental design and sample collection for scATAC-seq

Our initial objective was to investigate changes in the epigenetic landscape underlying the progression of the RGC developing trajectory, although information of other lineages was also obtained. Previous scRNA-seq analyses have demonstrated that there are four major cell states during RGC genesis, namely naïve RPCs (nRPCs), transitional RPCs (tRPCs), RGC precursors (early RGCs), and more differentiated RGCs (late RGCs), and each state is characterized by the expression of unique marker genes (22). Considering that cells in different states vary in numbers significantly, we enriched cells in individual states by fluorescence assisted cell sorting (FACS) utilizing two mouse lines we have generated, *Atoh7*^*zsGreen*^ (a null allele) and *Pou4f2*^*tdTomato*^ (a wild-type allele) (53). In mouse embryos carrying these alleles, tRPCs and fate-committed RGCs are marked with zsGreen and tdTomato respectively (**Figure 1A**) (53). Due to the stability of zsGreen, *Atoh7*^*zsGreen*^ also labels, and thereby allows for purification of, all early retinal cell types, including RGCs, H&As, and PHCs (mostly cones), as Atoh7 marks all tRPCs (22, 53). In this study, we collected a total of seven retinal cell samples, including double-negative cells from E14.5 *Atoh7*^*zsGreen/+*^;*Pou4f2*^*tdTomato/+*^ retinas (enriched with wild-type nRPCs and non-RGC lineage cells), zsGreen-expressing cells from E14.5 and E17.5 *Atoh7*^*zsGreen/+*^ retinas (considered as wild-type, thus enriched with wild-type tRPCs and their progenies), zsGreen-expressing cells from E14.5 and E17.5 *Atoh7*^*zsGreen/lacz*^ (*Atoh7*-null) retinas (enriched with *Atoh7*-null tRPCs and their progenies), and tdTomato-expressing cells from E14.5 and E17.5 *Pou4f2*^*tdTomato/+*^ cells (enriched with wild-type RGCs). The stability of the fluorescent proteins and the low gating threshold we purposely applied (**Suppl. Figure 1**) in FACS allowed us to isolate overlapping cell populations and achieve continuity along the developmental trajectories (53). 10X Chromium scATAC-seq libraries with these cell samples were generated and then sequenced. The sequence reads were then processed using Cell Ranger to identify insertion sites by Tn5 transposase in the genome in individual cells. After filtering out low-quality and doublet cells, we were able to obtain insertion data for 13,108 E14.5 cells and 8,910 E17.5 cells (**Suppl. Materials**). The quality of data for individual samples was assessed by enrichment in promoter regions and sizes of the excised DNA fragments, which were in good agreement with previous reports (54, 63) (**Suppl. Figure 2**). The Tn5 insertions per cell ranged from 24,963 to 43,867. We then used the MACS2 to identify reproducible peaks, defined as binned insertion sites per 500 bp window. A total of 261,595 reproducible peaks from the E14.5 samples and 251,029 reproducible peaks from the E17.5 samples were identified. These peaks were distributed at both gene bodies and intergenic regions and were likely located in regulatory elements such as active enhancers, poised enhancers, and promoters (loosely referred to as enhancers hereafter for simplicity) that were active at these two developmental stages (42, 63–65).

**Figure 1.**
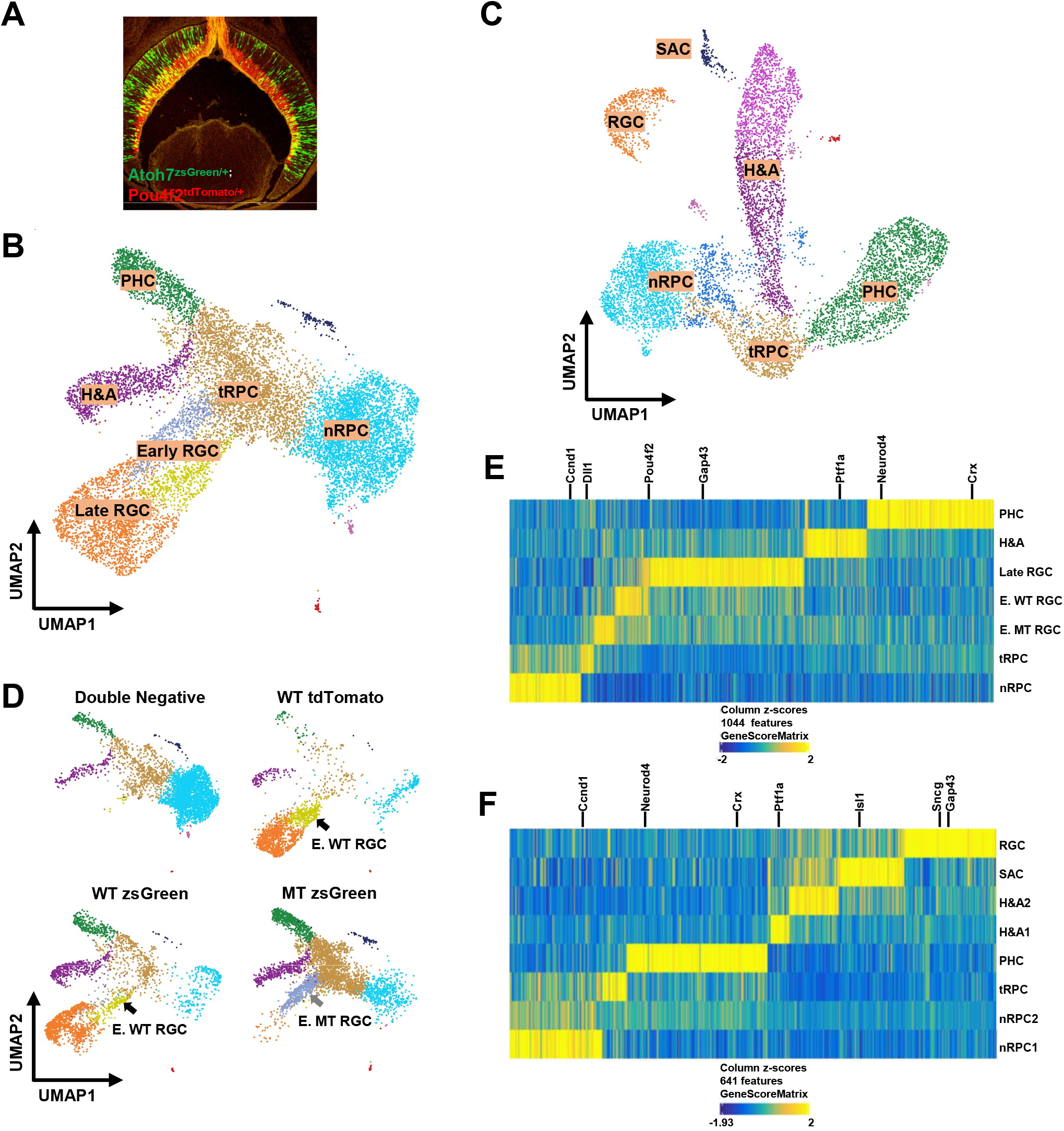
scATAC-seq of E14.5 and E17.5 retinal cells generates distinct clusters representing different cell states/types. **A**. zsGreen (green) and tdTomato (red) fluorescence from a retinal section from an *Atoh7*^*zsGreen/+*^*;Pou4f2*^*tdTomato/+*^ embryo demonstrating the two alleles mark two continuous stages of RGC differentiation. **B**. UMAP representation of different E14.5 clusters based on scATAC-seq. Identities of these clusters are marked based on marker gene activities as determined by GeneScore (see **E**). Note that there are two early RGC clusters, the dark blue one is made of Atoh7-null (MT) cells and the yellow one is made of wild-type (MT) cells (see **D**). **C**. UMAP representation of different E17.5 clusters based on scATAC-seq. Identities of these clusters are marked based on marker gene activities (see **F**). **D**. Contributions of the different FACS-purified E14.5 samples to individual clusters. The wild-type (WT) and *Atoh7*-null (MT) cells contribute to two distinct early RGC clusters as indicated by the arrows. **E. F**. Z score heatmaps based on gene activities to identify genes specifically activated in distinct E14.5 and E17.5 clusters. Selected marker genes for individual cell states/types are highlighted.

### scATAC-seq identifies the same cell states/types as scRNA-seq

Next, we used ArchR (54) to perform dimension-reduction clustering analysis of the E14.5 and E17.5 cells and displayed the results by Uniform Manifold Approximation and Projection (UMAP). The results allowed us to obtain 7 clusters from the E14.5 cells and 8 clusters from the E17.5 cells (**Figure 1B, C**). We then performed gene activity analysis using the GeneScore model, which predicts gene activities in scATAC-seq clusters with high accuracy (54), to identify genes with cluster-specific patterns. Unique sets of genes specifically active in each cluster were identified at the two stages (**Suppl. Dataset 1**), which could be represented by z-score based heatmaps (**Figure 1E, F**) and feature heatmaps (**Suppl. Figures 3, 4**). Examination of active genes in each cluster at the two stages revealed many known marker genes including *Ccnd1* and *Fgf15* for nRPCs, *Dll1* and *Gadd45a* for tRPCs, *Pou4f2, Isl1, Sncg* and *Gap43* for RGCs, *Ptf1a* for horizontal and amacrine cells (H&As), and *Crx* and *Neurod4* or photoreceptors (PHCs), (**Figure 1E, G**; **Suppl. Figures 3, 4, Suppl. Dataset 1**), and allowed us to unequivocally assign the identities of each cluster. The E14.5 clusters included nRPCs (1), tRPCs (1), H&As (1), PHCs (1), early RGCs (2, wild-type, MT, and mutant, WT), and late RGCs (1) (**Figure 1B, D, E**). The E17.5 clusters included nRPCs (2), tRPCs (1), H&As (2), PHCs (1), and RGCs (1) (**Figure 1C, F**). The differences between the two E17.5 nRPC clusters were not known, but they could be in different phases of the cell cycle or at different stages toward differentiation. There was one additional E17.5 cluster, which was likely starburst amacrine cells (SACs) since cells in this cluster expressed *Ptf1a, Sox2, Isl1, Chat*, and *Megf10*, which together mark SACs (66, 67) (**Figure 1C; Suppl. Figure 4, Suppl. Dataset 1**). A similar E17.5 cluster was observed by scRNA-seq previously (22).

As expected, cells from different FACS purified samples contributed differently to the different clusters and further supported the identity assignments to the clusters (**Figure 1D, Suppl. Figure 4**). Thus, double-negative cells from E14.5 *Atoh7*^*zsGreen/+*^;*Pou4f2*^*tdTomato/+*^ retinas contributed mostly to the nRPC cluster but not the RGC clusters. tdTomato+ cells from E14.5 *Pou4f2*^*tdTomato/+*^ retinas contributed mostly to RGC clusters. zsGreen positive cells from E14.5 *Atoh7*^*zsGreen/+*^ and *Atoh7*^*zsGreen/lacZ*^ retinas contributed mostly to tRPCs and their fate-committed progenies. Importantly, wild-type (WT) zsGreen positive cells from *Atoh7*^*zsGreen/+*^ retinas contributed to clusters representing all retinal cell types born at this stage, including RGCs (early and late), H&As, and PHCs (cones), but *Atoh7*-null (MT) zsGreen positive cells from *Atoh7*^*zsGreen/lacz*^ retinas contributed very few cells to the late RGC cluster, since RGCs die in the *Atoh7*-null retina (22, 31, 68) (**Figure 1D**). As expected, our low-threshold gating strategy (**Suppl. Figure 1**) in FACS resulted in overlapping cell populations from individual samples and enabled continuities of the developing trajectories (**Figure 1D**). In addition, the wild-type and *Atoh7*-null cells formed two neighboring early RGC clusters (**Figure 1B, D**), but such segregations did not occur in other clusters, indicating that the epigenetic differences between wild-type (WT) and *Atoh7*-null (MT) retinas existed mainly at this cell state. Similar but less obvious sample-specific contributions to the different clusters were observed in the E17.5 samples; noticeably, *Atoh7*-null cells did not contribute to the RGC cluster (**Suppl. Figure 4**). Although we did not specifically purify E17.5 double negative cells, a large number of nRPCs were included in the sorted cells and formed two clusters, likely due to the low gating cutoff threshold we used in FACS (**Suppl. Figures 1, 4**).

The cluster identities and their topographical relationships in the UMAP projections were in good agreement with those observed from our previous scRNA-seq clusters using cells from either the whole retina at E13.5 (22) or the same FACS purified cells at E14.5 (**Figure 1B, C, Suppl. Figure 5**) and E17.5 (22), suggesting that scATAC-seq data recapitulated the trajectories of the three major differentiation branches, including RGCs, H&As, and PHCs, during early retinal development (8, 21, 22). Unlike E14.5, the E17.5 RGC cluster was well separated from the other clusters and not connected with the tRPC cluster, and no E17.5 early RGC cluster was identified (**Figure 1C**). This was similarly observed with the E17.5 scRNA-seq (22), likely because at E17.5 RGC genesis was largely completed and very few tRPC were differentiating into the RGC lineage (6, 52). These findings indicated that the cell samples we collected adequately represented all cell populations present in the two developmental stages.

### Differentially accessible peaks reveal the shifting epigenetic landscape along individual developmental trajectories

The different scATAC-seq clusters were defined by the differentially accessible peaks in them, which represented potential cell state/type specific enhancers. A total of 127,302 differentially accessible peaks in the E14.5 samples and 151,719 in the E17.5 samples were identified, and unique sets of differentially accessible peaks were identified for each E14.5 and E17.5 cluster (**Suppl. Datasets 2, 3**). Z score clustering based on these peaks further validated the relationships among these clusters at both E14.5 and E17.5, following the developmental trajectories from nRPCs to tRPCs, and then to the different cell fates including H&As, PHCs, and RGCs (**Figure 2A**).

**Figure 2.**
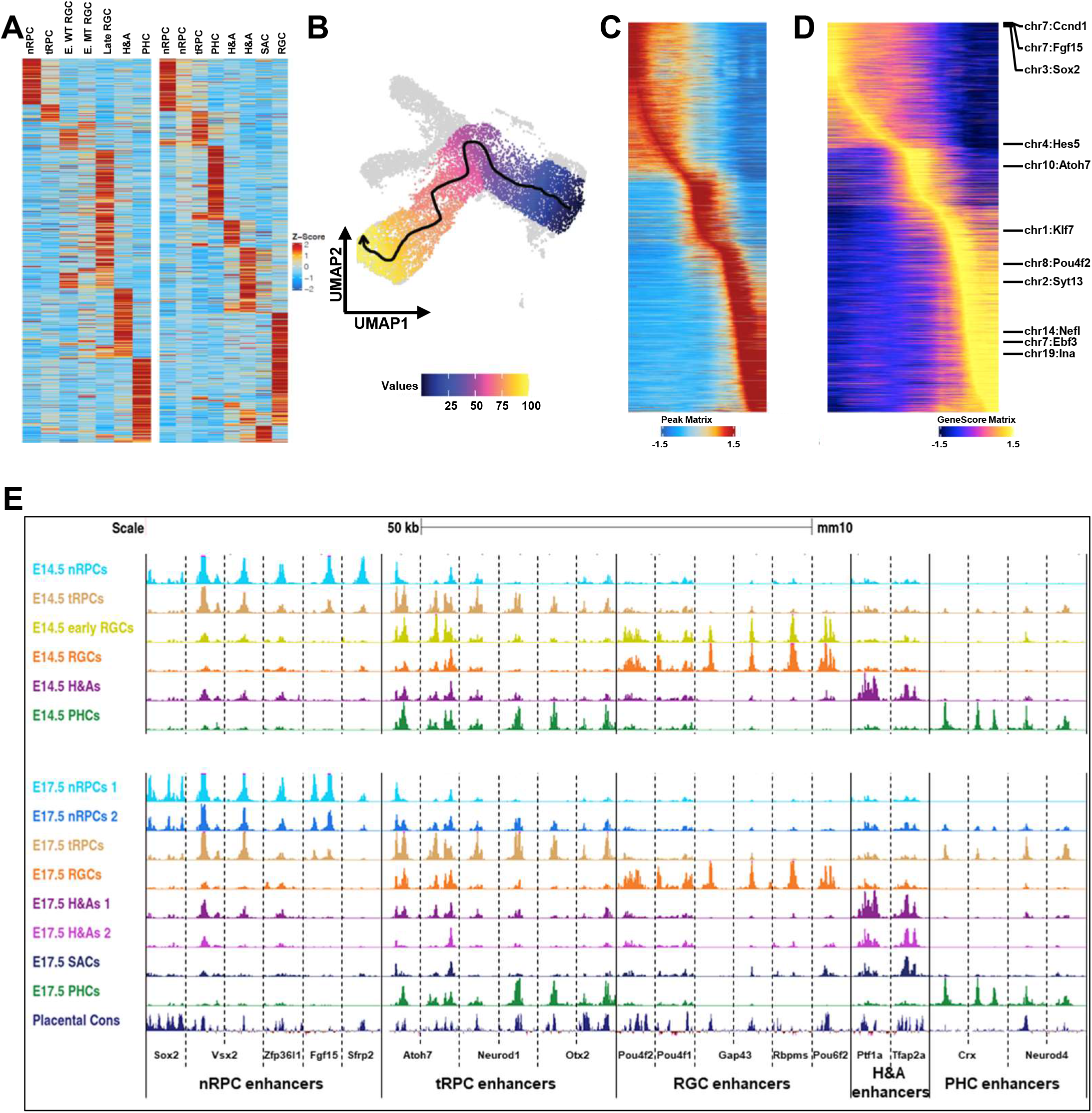
The shifting epigenetic landscape is defined by retinal cell state/type specific enhancers. **A**. Z score heatmaps displaying the unique sets accessible peaks identified for each cell state/type at both E14.5 (left panel) and E17.5 (right panel). **B**. Identification of the developmental trajectory of the RGC lineage from nRPCs to late RGCs (arrow indicates the direction of the trajectory). **C**. The shifting epigenetic landscape along the developmental trajectory of the RGC lineage (from left to right). Each horizontal line represents a differentially accessible peak. **D**. Shifting of gene activity along the developmental trajectory of the RGC lineage. Examples of known marker genes expressed in different cell states/types are highlighted. **E**. Enhancers associated with genes expressed in specific cell states/types display distinct dynamics along individual developmental trajectories. scATAC-seq tracks of selected regions for individual cell states/types at E14.5 and E17.5 are displayed (genome coordinates (mm10) of these regions are provided in **Suppl. Materials**). The bottom track displays the conservation of these enhancers among placental mammals.

The shifting epigenetic landscape was further confirmed and illustrated by trajectory analysis for the RGC lineage (54), which clearly identified its direction from nRPCs to tRPCs, then to early RGCs, and finally to more differentiated (late) RGCs (**Figure 2B, C**). Noticeably, the trajectory progressed from *Atoh7*-null (MT) early RGCs to wild-type early RGCs (**Figures 1D, 2B**), indicating that Atoh7 was required for the normal progression. The changes of accessibilities of the differentially accessible peaks along this trajectory not only confirmed their cluster specificities but also their changes from one state (cluster) to the next, exhibiting the shifting of the epigenetic landscape as differentiation progressed (**Figure 2C**). In addition, the gene activities of the cluster-specific genes, including many known marker genes, as determined by GeneScore, along the trajectory displayed very similar shifts (**Figure 2D**). Since the GeneScore activities for the marker genes were consistent with the actual gene expression patterns revealed by scRNA-seq (22), the shifting epigenetic landscape along individual developmental trajectories, including that of the RGC lineage, likely underlay the corresponding changes in gene expression.

The shifts of the epigenetic landscape were further confirmed by examining enhancers associated with selected cell state/type-specific genes such as *Sox2, Vsx2, Zfp36l1, Fgf15*, and *Sfrp2* for nRPCs, *Atoh7, Otx2*, and *Neurod1*, for tRPCs, *Pou4f2, Pou4f1, Gap43, Rbpms* and *Pou6f2* for RGCs, *Crx* and *Neurod4* for PHCs, and *Ptf1a* and *Tfap2b* for H&As. Cell state/type-specific enhancers with activity dynamics closely mirroring that of the expression of these genes as revealed previously by scRNA-seq could be identified in their proximity at both E14.5 and E17.5 (22) (**Figure 2E**). Thus, enhancers associated with nRPC genes were most active (as judged by normalized Tn5 insertion frequencies) in nRPCs, less so in tRPCs, and largely inactivated in the different differentiated neurons. tRPC enhancers were often activated in nRPCs already, but only became most active in tRPCs, and, depending on what lineages the genes were involved in, remained active only in the relevant cell types; as such, enhancers associated with *Atoh7* remained active in multiple lineages including early RGCs, whereas those associated with *Neurod1* and *Otx2* remained active only in the PHCs. On the other hand, enhancers associated with genes expressed in differentiated neurons including RGCs, H&As, and PHCs only became active in the respective cell types when the cell fates were determined. Most of these enhancers were highly conserved, further supporting their functional significance (**Figure 2E**).

We were also interested in the temporal changes of the epigenetic landscape from E14.5 to E17.5. For that purpose, we pooled all the wild-type scATAC-seq data from the two stages together and performed clustering and GeneScore analysis. We obtained very similar clustering results as those obtained when the two stages were analyzed separately (**Suppl. Figure 6**). Identical cell states/types were identified and their relationships remained the same. Noticeably, except for SACs which were present only at E17.5, clusters representing corresponding cell states/types from the two stages were always close to each other but separated (**Suppl. Figure 6**). These could not have been caused by batch effect as the separations followed strictly by stages, but not samples. These observations indicated the epigenetic landscape shifted for all the cell states/types. We thus identified the differentially accessible peaks and associated genes based on gene scores by pairwise comparison of corresponding clusters of the two stages (**Suppl. Datasets 4, 5**). These peaks (enhancers) and associated genes likely reflected the changes of competence in the RPCs, shift of cell types being produced, and maturation of neurons generated earlier. Examples of such changes including upregulation of *Nfix* in nRPCs (**Suppl. Dataset 5**) and upregulation of *Otx2, Crx*, and *Neurod4* in tRPCs (**Suppl. Dataset 5**). This was consistent with the roles of Nfi factors in the shift of RPC competence to produce late retinal cell types (8, 69) and upregulation of *Otx2, Crx*, and *Neurod4* in tRPCs reflected the shift (15). More in-depth studies will be required to understand the process, but our data provide a very useful start for such studies.

### Differentially active enhancers regulate cell state/type specific genes

Next, we sought to link individual enhancers with the genes they likely regulate at the genome level. For that purpose, we performed data integration of the scRNA-seq data with the scATAC-seq data obtained with the same cell populations isolated by FACS at E14.5 (**Suppl. Figure 5**) and E17.5(22). This integration links cell-state/type specific enhancers with genes they likely regulate (peak to gene link, P2G link) based on the similarity of their dynamics across the different clusters and their proximities (54). For the E14.5 data set, we identified 55,293 P2G links to 6,600 genes, with the peaks and genes assigned to specific clusters (**Suppl. Dataset 2**). Similarly, for the E17.5 data, we identified 76,079 links to 8,531 genes (**Suppl. Dataset 3**). Often peaks from separated regions were assigned to a single gene, indicating that multiple enhancers were involved in regulating the gene. The accessibilities of the peaks, or activities of the corresponding enhancers, and the expression levels of genes they were linked to were highly correlated, as indicated by the heatmaps of the two sets of data for both E14.5 and E17.5 clustered by z scores (**Figure 3A, B**). We then compared the E14.5 and E17.5 linked gene sets with the 1,829 cell state/types enriched genes we previously identified by scRNA-seq using cells from the whole retina (22), and found 84.5% (1,545) of those genes were included in the E14.5 gene set and 90.3% (1,651) of those genes were included in the E17.5 gene set (**Figure 3C**). Thus, the P2G analysis allowed for the identification of putative enhancers and the genes they likely regulate and assessment of their activities in all the cell states/types at these two developmental stages.

**Figure 3.**
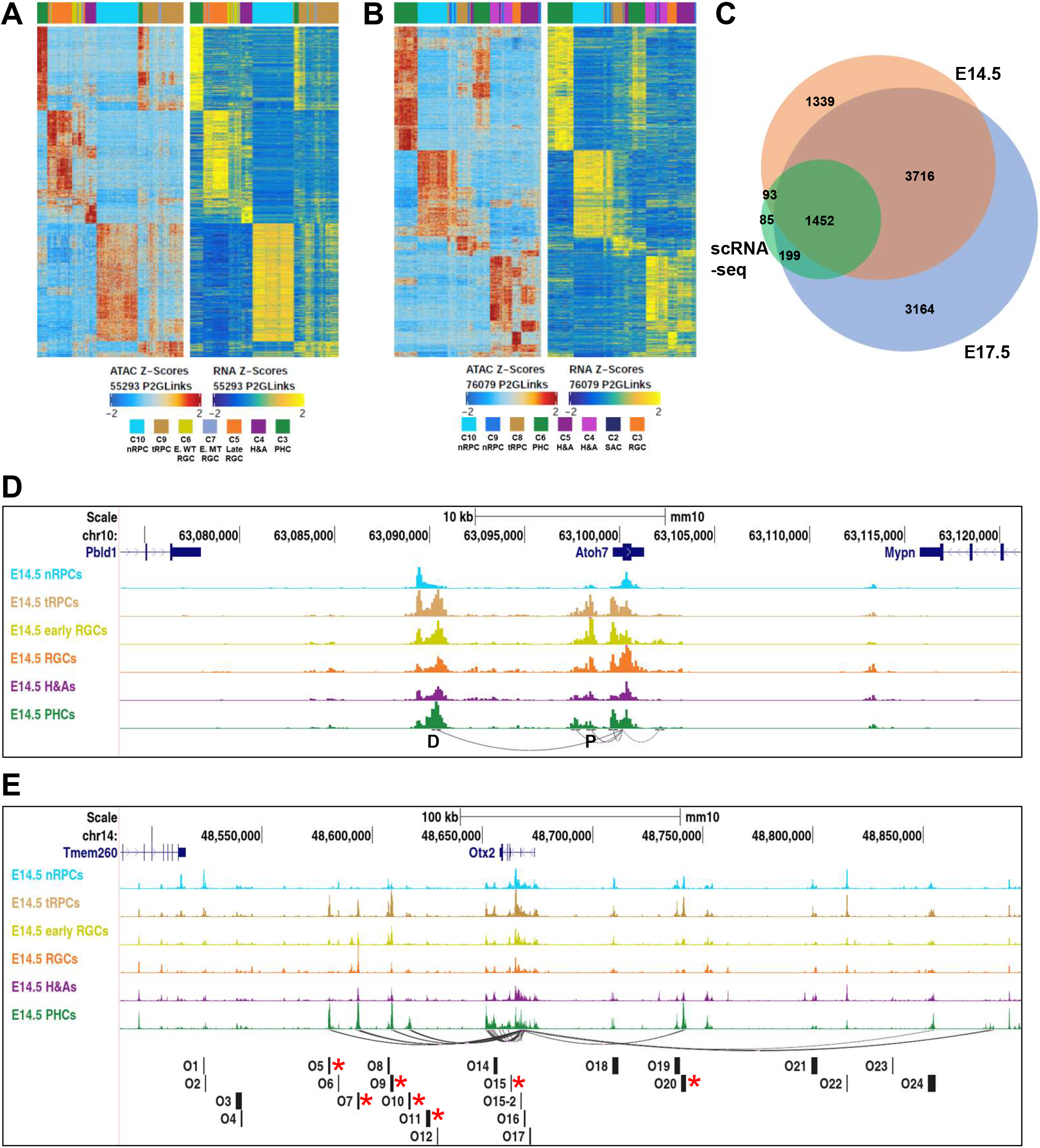
Peak to gene (P2G) linkage analysis identifies enhancers for individual genes globally. **A. B**. Heatmaps of linked scATAC-seq peaks and scRNA-seq genes at E14.5 and E17.5 respectively. **C**. A Venn diagram depicting the overlaps of P2G linked genes and the cell state/type-specific genes identified by scRNA-seq. **D**. Genome tracks displaying P2G linked enhancers for *Atoh7* and their dynamics across the different cell states/types. Arched lines denote P2G links by connecting the TSS to the enhancers. D indicates the distal enhancer and P indicates the proximal enhancer. **E**. Genome tracks displaying P2G linked enhancers for *Otx2* and their dynamics across the different cell states/types. The bottom track shows the positions of the candidate enhancers studied by Chan et al (2020); the seven active enhancers reported in that study are marked by red asterisks, six of which are identified by P2G but O11 is not active at this stage.

The validity of these candidate enhancers was further assessed by compassion with previously studied enhancers in *Atoh7* and *Otx2*, two genes both expressed in tRPCs but involved in two distinct lineages, RGCs and PHCs, respectively (29, 30, 34, 70). Two enhancers associated with *Atoh7* were previously reported, a proximal enhancer and a distal shadow enhancer (71–74). The P2G link analysis not only correctly identified the two enhancers but also revealed the dynamics in their accessibilities (activities) in the different cell states/types (**Figure 3D**). The activity dynamics of proximal enhancer (P) almost completely reflected that of *Atoh7* expression, silent in nRPCs, most active in tRPCs, and continued in the early RGCs, but largely diminished in late RGCs and other differentiated neurons. In contrast, the distal enhancer (D) contained two regions with distinct activity dynamics. The more distal part was active in nRPCs already, reached peak activity in tRGCs, but tapered off markedly in differentiated neurons including RGCs. The proximal part, on the other hand, was inactive in nRPCs, became most active in tRPCs, but remained active in many of the differentiated neurons. These enhancers may play different roles in the different phases of **Atoh7** activation and function collaboratively and sequentially to ensure normal expression. For *Otx2*, the P2G analysis recognized a set of enhancers that were predominantly active in tRPCs and/or PHCs (**Figure 3E**), many of which have been previously identified and studied (49, 75–78). Importantly, these P2G enhancers included six of the seven enhancers validated experimentally in a previous study that screened 22 potential enhancers (75), all with dynamics closely mirroring that of *Otx2* expression (**Figure 3E**). These results indicated the cell state/type specific enhancers were the major drivers for cell state/type specific expression.

Several general features were observed regarding the enhancers associated with cell state/type specific genes. Typically, as observed with *Atoh7* and *Otx2*, multiple state/type-specific enhancers were associated with individual genes. Many enhancers were often active in multiple continuous cell states along individual lineage trajectories (e.g. O5, O9, and O20 of *Otx2*, **Figure 3E**), reflecting the expression dynamics of the associated genes. Whereas most enhancers were found in the gene body and the immediate intergenic regions, some putative enhancers were found within a neighboring gene body or even over a neighboring gene (see later results). There were also enhancers being shared by neighboring genes, suggesting that the genes were co-regulated. Often genes with shared enhancers belonged to members of a gene family closely located on the chromosome (e.g. the *Irx* genes, *Nefl* and *Nefm*) (**Suppl. Figure 7A**). For genes expressed in multiple lineages, e.g., *Onecut1* (62, 79, 80), enhancers for the separate lineages existed (**Suppl. Figure 7B**). On the other hand, there were also cases in which cell state/type specific genes had no apparent corresponding enhancers with similar dynamics associated with them. This was particularly true for many genes involved in cell proliferation and expressed in nRPCs (22), including *Pcna* and the *Mcm* genes (*Mcm2-10*), although constitutively accessible enhancers were often present (**Suppl. Figure 7C-E**). It is possible that these genes were regulated by long distance enhancers, which were not readily identified. Alternatively, the constitutively accessible enhancers may be active only in nRPCs due to differential transcription factor binding.

### Transcription factors functioning in different cell states/types

The functions of enhancers are mediated by transcription factors bound to specific DNA sequences in them. Many transcription factors expressed and functioning in different stages of retinal cell differentiation have been identified (13–16). To examine whether the cell state/type specific enhancers we identified interacted with these transcription factors, we examined the enrichment of DNA motifs in the differentially accessible peaks by chromVAR (57). Unique sets of DNA motifs were enriched in each cluster at E14.5 and E17.5 (**Figure 4A, B, Suppl. Datasets 2, 3**), and many of these motifs were recognized by transcription factors known to function in the corresponding cellular states/types. Motifs enriched in distinct E14.5 clusters included those recognized by Vsx2, Rax, Lhx2, Pax6, Sox2 in nRPCs, bHLH transcription factors such as Atoh1/7, Neurod1, Ascl1, and Neurog2, and Otx2 in tRPCs, Otx2, Crx, and Onecut (Onecut1/2) in PHCs, Tfap2a and Onecut in H&As, and Pou4f2, Ebf (Ebf1-4), and Onecut in RGCs (**Figure 4A**). Interestingly the DNA motif bound by the Tead family (Tead1-4), which is part of the Hippo pathway (81, 82), was ranked top in nRPCs, indicating that the Hippo pathway likely played important roles in nRPCs, although this needs to be further investigated. Neighboring cell states tended to have shared enriched motifs, further highlighting the continuation and transition from one state to the next along the developmental trajectories of the different lineages. The best example of such transition was the tRPCs, which not only had enriched motifs for the nRPCs, albeit at lower levels, but also those recognized by transcription factors functioning in differentiated neurons. There were also cases where chromVAR identified motifs associated with transcription factors not aligned with their known expression patterns and functions, such as motifs for Isl1 and Myt1l (**Figure 4A**), which are expressed in RGCs (39, 83) but chromVAR identified their binding sites enriched in nRPCs. This likely was due to limitations of the transcription factor motif databases available. The E17.5 clusters share many top ranked motifs with E14.5 as revealed by chromVAR (**Figure 4B**), but there were noticeable differences. For example, the Onecut motif was not enriched in any clusters at E17.5 These differences likely reflected changes in the epigenetic landscape associated with the temporal progression of retinal development.

**Figure 4.**
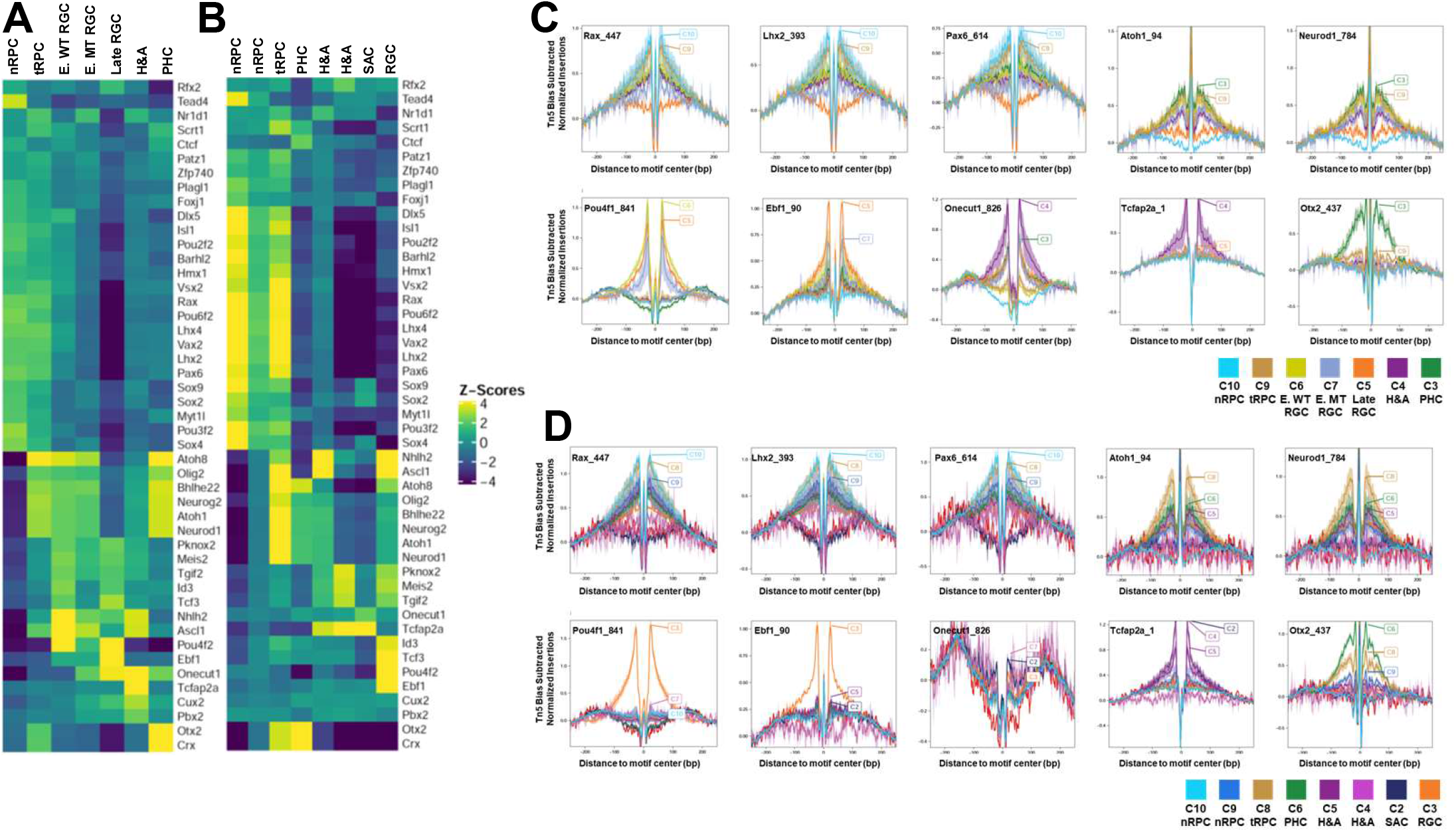
DNA motif enrichment and footprinting analysis reveal cell state/type specific occupation of key transcription factors. **A.B**. chromVAR analysis of DNA motifs recognized by distinct transcription factors enriched in scATAC-seq peaks in a cell state/type specific manner at E14.5 and E17.5. **C.D**. Cell state/type-specific footprinting at E14.5 and E17.5 respectively of selected transcription factors as revealed by reduction in Tn5 insertion frequencies around the relevant DNA motifs. Y-axis represents normalized insertion frequencies with the Tn5 incision bias subtracted.

To further validate that the above indicated transcription factors indeed bound enhancers in a cell state/type specific fashion, we also performed footprinting analysis based on the scATAC-seq data (54), and observed cell state/type specific footprints for many of these sequence motifs. At E14.5, footprints were found for Lhx2, Rax, and Pax6 in nRPCs and tRPCs, for Neurod1 and Atoh7 (Atoh1) in tRPCs and PHCs, for Pou4f (Pou4f1-3) in early and late RGCs, for Ebf (1-4) factors in late RGCs, for Tfap2a in H&As, for Onecut1/2 in RGCs, H&As, and PHCs, and for Otx2 in tRPCs and PHCs (**Figure 4C**). Similar cluster-specific footprints were also observed at E17.5 (**Figure 4D**). A comparison of footprints at E14.5 and E17.5 further confirmed the temporal shifts of the epigenetic landscape. As such, at both E14.5 and E17.5, clear Pou4f (1-3) footprints were seen in clusters of the RGC lineage (**Figure 4C, D**), indicating that Pou4f factors continue to function in RGCs even after RGC genesis had largely completed. In contrast, footprints for the Onecut factors were prominent at E14.5 in all three lineages, but they were largely diminished in these lineages at E17.5 (**Figure 4D**), an observation consistent with the results from the motif enrichment analysis (**Figure 4A, B**) as well as their reduced expression levels at later stages (80).

These findings suggest that key transcription factors indeed bind to the differentially active enhancers in specific cell states/types along the developmental trajectories, and they likely are involved not only in shaping the epigenetic landscape in a particular state but also in driving it to the next state.

### The epigenetic landscape in transitional RPCs

One of the key findings by scRNA-seq was that all retinal trajectories go through a shared state, which were called neurogenic or transitional RPCs (8, 21, 22). The tRPCs are characterized by the expression of genes encoding components of the Notch pathway, including *Dll1, Dll3, Dll4, Notch1, Hes5, Mfng*, and *Rpbj* as well as co-expression of genes encoding transcription factors for distinct lineages (22). The underlying epigenetic landscape in these cells should provide insights into how this cell state is established and maintained, and how it diverges into the different retinal lineages. tRPCs were characterized by a unique set of active enhancers, which likely underlay the expression of genes in this cell state (**Figure 2A**). Consistently, the P2G analysis identified tRPC-specific enhancers for all the lineage specific genes, including *Atoh7, Sox4, Sox11, NeuroD1, Otx2, Onecut1, Neurog2, Olig2, Foxn4*, and the Notch pathway genes (examples shown in **Figure 2E, Figure 3D, E, Suppl. Figures 7B, 8, 10**). Some of these enhancers were active in nRPCs already, whereas others remained active in the more differentiated stages, including PHCs and RGCs (**Figure 2E, Figure 3D, E, Suppl. Figures 7B, 8, 10**). These observations underscored the transitional nature of this cell state.

Compared to nRPCs, the most prominent change in the epigenetic status of tRPCs was the drastic enrichment of DNA motifs recognized by bHLH transcription factors and Otx2 (**Figure 4A, B**). In the early embryonic retina, multiple bHLH transcription factors, including Atoh7, Neurod1, Neurog2, Olig2, Ptf1a, Bhlhe22, and Ascl1, which all binds to the E box motif, are expressed in tRPCs. These bHLH factors all have distinct roles in retinal cell differentiation (18, 84–90). Consistent with these bHLH factors functioning in retinal cell differentiation, the E box motifs were not enriched in the nRPCs, but continued to be enriched in early RGCs (but not late RGCs), H&As, and PHCs (**Figure 4A, B**). These findings indicated that activation of these bHLH transcription factor genes and their impact on the tRPC epigenetic landscape, e.g., activation of enhancers containing E boxes, likely were the first steps of retinal cell differentiation. Other transcription factors such as Otx2 likely also play critical roles in initiating retinal cell differentiation. Consistently, the Otx2 DNA motif was also highly enriched in tRPCs, and remained so in PHCs but not other differentiated neurons (**Figure 4A, B**).

Since Atoh7 and Otx2 drive tRPCs to two different cell fates (29, 30, 70), learning how they interact with and influence the tRPC epigenetic landscape may shed light on how the two corresponding lineages emerge. To that end, we performed CUT&Tag (58) to identify the genomic sites bound by Atoh7 and Otx2 in the E14.5 retina. Independent duplicate experiments for each transcription factor were performed and high reproducibility (correlation coefficients 0.845 and 0.946 for Atoh7 and Otx2 respectively) was observed. We then used the merged sequence reads to call for peaks for both Atoh7 and Otx2 by MACS2 (56) and further filtered these peaks by intersecting them with the E14.5 enhancers identified by scATAC-seq, which allowed us to identify active enhancers containing binding sites for Atoh7 and Otx2 respectively. A total of 6,626 Atoh7-bound and 4,992 Otx2-bound active enhancers were identified (**Suppl. Dataset 6**); these enhancers likely mediate the roles of the two transcription factors. Importantly, the activities of these enhancers, as determined by the normalized Tn5 insertion frequency in the scATAC-seq data, were consistent with the roles of these two factors in retinal cell differentiation (14, 15, 22, 70). Enhancers bound by Atoh7 were most active in tRPCs and early RGCs, whereas enhancers bound by Otx2 were most active in both tRPCs and PHCs or just PHCs, as visualized by heatmaps **(Figure 5A, B**) and genome tracks of example enhancers **(Figure 5G**). HOMER motif enrichment analysis (91) on these enhancers found that the top enriched motifs were those recognized by the two transcription factors respectively; 69.30% of the Atoh7 enhancers contained the E box motif recognized by Atoh1/7, whereas 70.96% of the Otx2 enhancers contained the Otx2/Crx motif **(Figure 5D, E**), validating that CUT&Tag indeed identified bona fide binding sites for these two transcription factors. The activities of the enhancers interacting with Atoh7 and Otx2 further established that they both function in tRPCs and the respective developing neurons, RGCs and PHCs.

**Figure 5.**
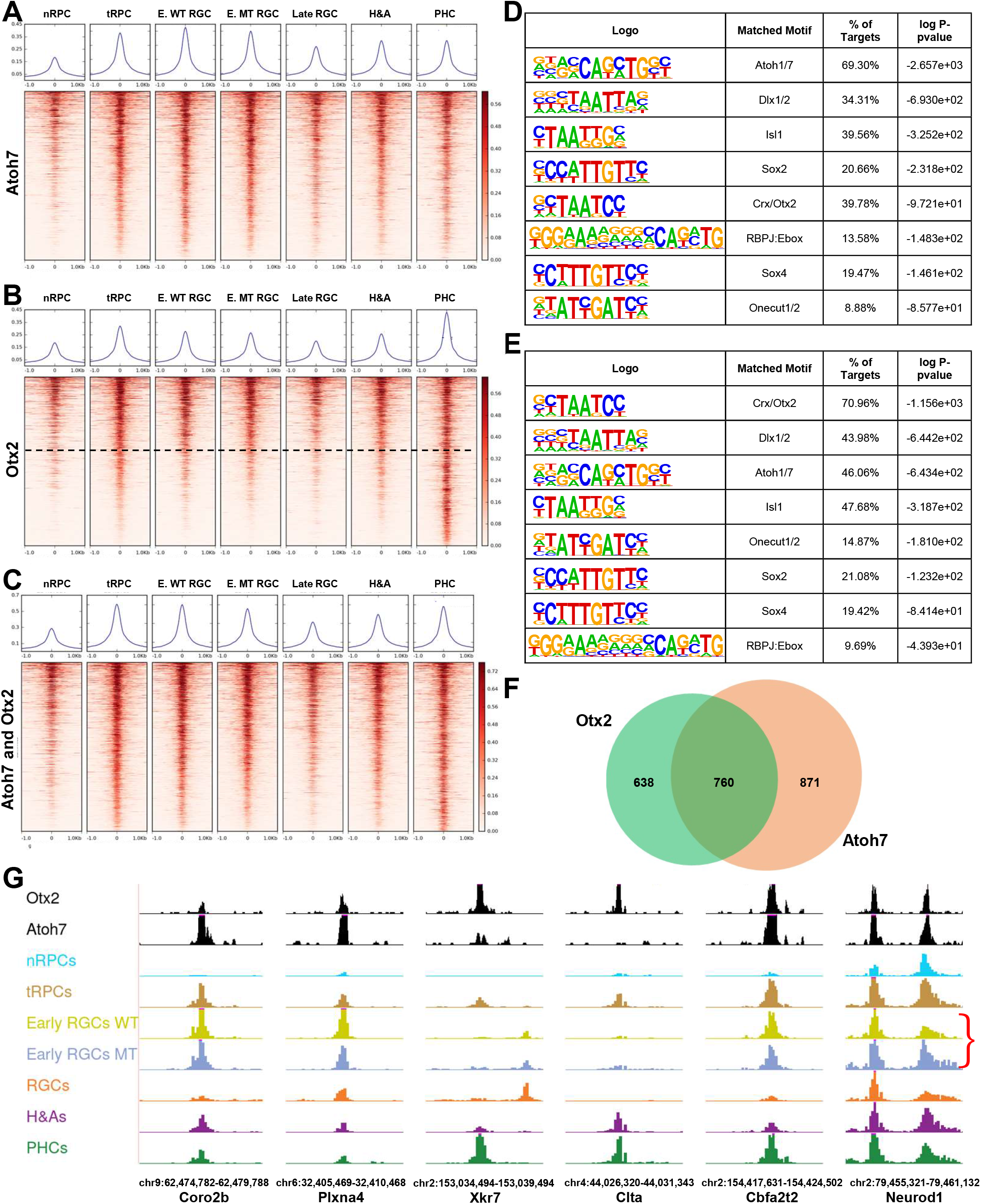
Atoh7 and Otx2 bind to distinct but overlapping sets of enhancers. **A**. Activities of Atoh7-bound enhancers as determined by normalized Tn5 insertion frequencies across cell states/types at E14.5. **B**. Activities of Otx2-bound enhancers as determined by normalized Tn5 insertion frequencies across cell states/types at E14.5. The horizontal dashed line demarcates enhancers with similar activities in tRPCs and PHCs from those more active in PHCs. **C**. Activities of enhancers co-bound by Atoh7 and Otx2 across cell states/types. For **A-C**, genomic regions of the MACS2 summits ±1.0 kb were displayed. In **C** the Atoh7 summits were used as centers to generate the heatmap. The curves on top of each heatmap show the average Tn5 insertion frequencies of all peaks in individual cell states/types. **D**. Motif enrichment analysis by HOMER of Atoh7-bound enhancers displayed in **A. E**. Motif enrichment analysis by HOMER of Otx2-bound enhancers displayed in **B. F**. A Venn diagram depicting the overlap of genes likely regulated by Atoh7 and Otx2. **G**. Example enhancers bound by Atoh7 and/or Otx2 and their cell state/type-specific activities as displayed in the genome browser. The red bracket in the last panel indicates the upregulation of one of the *Neurod1* enhancers in the *Atoh7*-null retina. Genome coordinates (mm10) of the displayed regions and the nearby genes are indicated at the bottom.

### Atoh7 and Otx2 cross-regulate each other and downstream genes to drive tRPCs toward two distinct lineages

Additional motifs for transcription factors known to function in retinal cell differentiation were enriched in both the Atoh7- and Otx2-bound enhancers (**Figure 5D, E**), indicating that the two factors collaborate with other transcription factors to regulate downstream genes. Such transcription factors included those functioning in nRPCs such as Sox2, in tRPCs such as the SoxC factors (e.g. Sox4) and the Onecut factors, and in differentiated neurons such as Dlx1/2 and Isl1, further highlighting the transitional nature of this cell state. In addition, a composite motif containing the binding site for Rbpj, an effector of the Notch pathway, and an E box (RBPJ:Ebox motif) was also significantly enriched in enhancers bound by Atoh7 and/or Otx2 (**Figure 5D, E**). The RBPJ:Ebox motif, which regulates the expression of bHLH proneural genes, such as *Neurog2* in the spinal cord and *Atoh7* in the retina (92, 93), likely mediates the functions of the Notch pathway in establishing the tRPC state. Of notice was that the Otx2/Crx motif was highly enriched (39.78%) in the Atoh7 enhancers, and vice versa, the Atoh1/7 E box motif was also highly enriched in the Otx2 enhancers (46.06%) (**Figure 5D, E**). In agreement, we found that there was a substantial overlap of the Atoh7 and Otx2 enhancer sets; 2,059 enhancers were co-bound by Atoh7 and Otx2 (**Figure 5C, Suppl. Dataset 7)**. These enhancers were generally more active in tRPCs, early RGCs and PHCs than in other cell states/types (**Figure 5C, G, Suppl. Dataset 7**). Noticeably, these enhancers did not include those that were most active in PHCs (note the lower part of **Figure 5B**). These observations indicated that Atoh7 and Otx2 interact and co-regulate target genes at the early stage of differentiation, mostly in tRPCs, but their functions diverge as tRPCs differentiate into distinct cell types.

Based on the P2G data, we were able to identify genes associated with the Atoh7 and Otx2 enhancers, 1,631 for Atoh7 and 1,398 for Otx2. As expected, Atoh7 target genes included those functioning in RGC differentiation and function, e.g., *Pou4f2* and *Isl1* (**Suppl. Dataset 6**), whereas Otx2 target genes contained those involved in photoreceptor differentiation and function, e.g., *Neurod1, Neurod4*, and *Crx* (**Suppl. Dataset 6**). On the other hand, and consistent with the two factors co-binding to many enhancers, many (760) genes were targets of both Atoh7 and Otx2 (**Suppl. Dataset 7, Figure 5F**). These genes comprised the many key regulatory genes expressed in tRPCs such as *Atoh7* and *Otx2* themselves and *Foxn4, Olig2, Neurod1, Ascl1, Onecut1, Insm1*, and *Dlx1/2* (**Suppl. Dataset 7, Figure 6**), downstream regulatory genes functioning in differentiating RGCs (e.g. *Pou4f2, Isl1*, and *Klf7*) or PHCs (e.g. *Neruod4*) (**Suppl. Dataset 7, Figure 6, Figure 9)**, and component genes of the Notch pathway, including *Dll1, Dll4, Mfng, Notch1, Hes1, Hes5*, and *Hes6* (**Suppl. Dataset 7, Figure 6**). As noted earlier, for individual genes, multiple enhancers were often involved, some bound by both factors, whereas others by only one of them (**Figure 6**).

**Figure 6.**
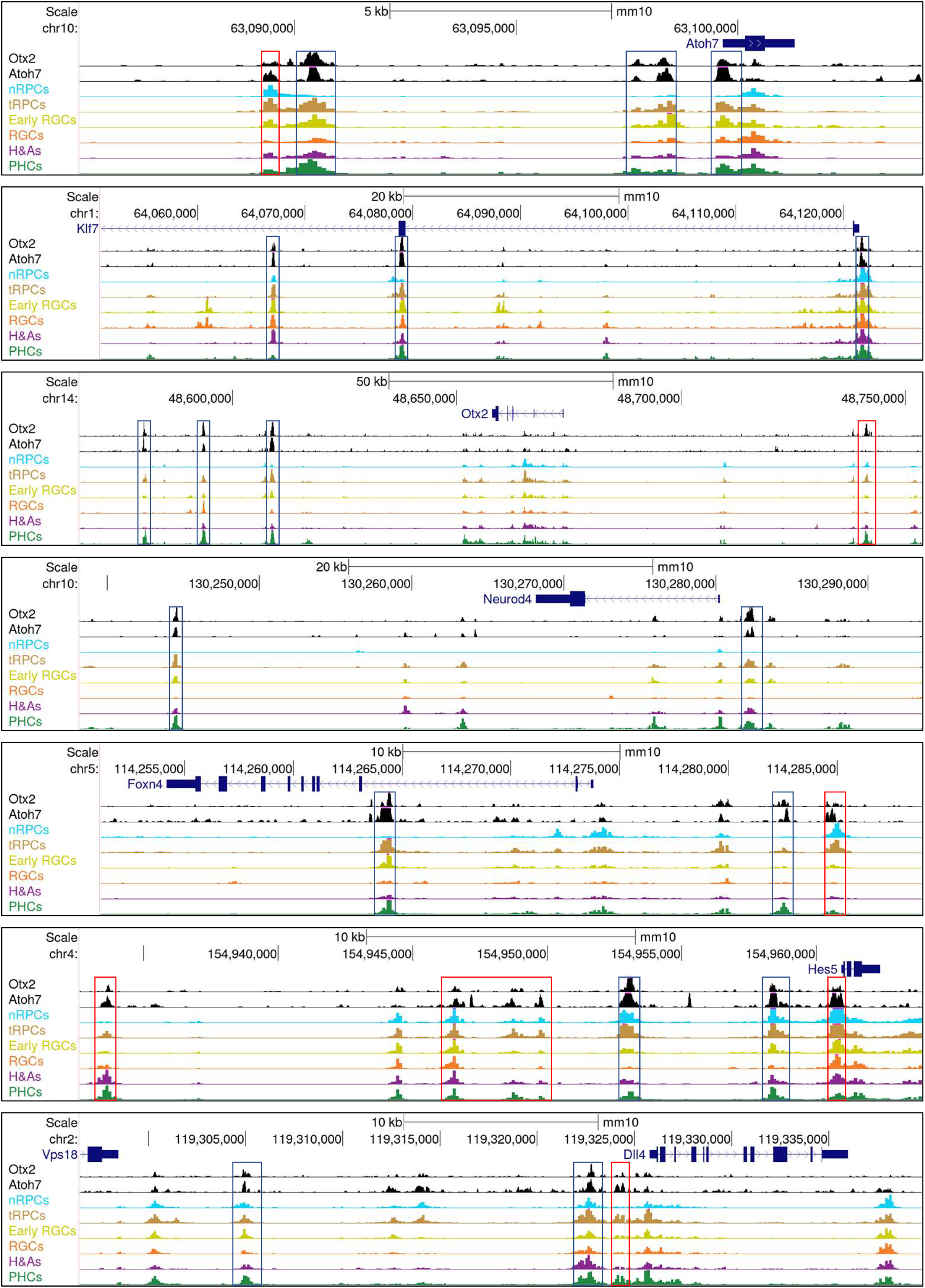
Example genes are co-regulated by Atoh7 and Otx2. Displayed are Atoh7 and Otx2 CUT&Tag genome tracks as well as the E14.5 scATAC-seq tracks of regions surrounding genes involved in the formation of the RGC (*Atoh7* and *Klf7*), PHC (*Otx2* and *Neurod4*), and H&A (*Foxn4*) lineages, as well as two genes of the Notch pathway (*Hes5* and *Dll4*). Note that for each gene, there are multiple enhancers co-bound by Atoh7 and Otx2 (blue boxes), although there are also enhancers predominately bound by just one of them (red boxes).

The respective contributions by Atoh7 and Otx2 to the final expression of their target genes are likely gene and cell context specific. In some cases, such as the Notch pathway genes, they may cooperate to activate the target genes. Many of the Notch pathway genes have been reported to be dependent on Atoh7 (22, 38, 94, 95). Our results indicate that Atoh7 regulates these genes directly and that Otx2 may collaborate with Atoh7 in this process. In the cases of the lineage specific genes, they may function in opposite directions, as they promote two distinct lineages. Thus, in the latter scenario, Atoh7 promotes RGC genes such as *Atoh7, Pou4f2*, and *Isl1* and represses PHC genes such *Otx2, Neurod1, Neurod4*, and *Crx*, whereas Otx2 does the opposite by promoting the PHC genes but repressing the RGC genes including *Atoh7*. Supporting this idea is their co-regulation of *Neurod1*, which is activated by Otx2 but repressed by Atoh7 (22, 38, 96). This is likely mediated by multiple *Neurod1* enhancers bound by both Atoh7 and Otx2, and the repressive role of Atoh7 was supported by the increased activities of these enhancers in the *Atoh7*-null retinas (**Figure 5G**). Since both Aoth7 and Otx2 also bind to enhancers associated with H&A genes such as *Foxn4* (**Figure 6**), the cross repression of the lineage specific genetic programs by key regulators is likely a general mechanism by which individual cell fates arise from the multipotent tRPCs.

### Atoh7, Pou4f2 and Isl1 interact with the epigenome to activate RGC-specific genes in a sequential and combinatorial fashion

The establishment of the RGC lineage involves a cascade of gene regulation by state-specific transcription factors, including Atoh7 in tRPCs and early RGCs, and Pou4f2, Isl1, and many other transcription factors in early and late RGCs (14). When the overall activities of enhancers bound by Atoh7 were compared between wild-type and *Atoh7*-null retinas, only moderate reductions were observed in the early RGC state (**Figure 5A**). To further examine how Atoh7 impacted the epigenetic landscape, we compared corresponding E14.5 wild-type and *Atoh7*-null clusters to identify the enhancers dependent on Atoh7 and their associated genes. We identified very few differentially accessible peaks in nRPCs, H&As, and PHCs (**Suppl. Dataset 8**). Of notice was that enhancers associated with two sonic hedgehog (Shh) pathway genes, *Gli1* and *Ptch1*, were highly reduced in activity in the *Atoh7*-null nRPCs. This was consistent with this pathway being compromised in the Atoh7-null retina as RGCs, which are the source of Shh, are absent (37, 38, 97, 98). However, we found 473 scATAC-seq peaks in tRPCs that were dependent on Atoh7 (**Suppl. Dataset 8**). In early RGCs, we identified 11,532 Atoh7-dependent peaks (**Suppl. Dataset 8**). These results indicated that Atoh7 regulates the activities of just a specific subset of enhancers. Moreover, when we separated the WT and MT scATAC-seq and scRNA-seq data and performed P2G link analysis on this set of enhancers, we identified 2,647 links and observed a very high degree of correlation between epigenetic and expression changes as demonstrated by the z score maps (**Suppl. Figure 9**). Genes associated with Atoh7-activated enhancers included a large number of key regulatory genes involved in the RGC lineage, such as *Isl1, Pou4f2, Pou4f1, Eya2, Klf7, Sox11, Irx2, Dlx2*, Lmo1, *Myt1l, Lmo1*, and *Tle1*, and some RGC structural/functional genes such as *Elavl4, Dcc, Vav3*, and *Stmn2* (**Suppl. Dataset 8**). In contrast, and as mentioned above, the enhancers repressed by Atoh7 were associated with upregulated genes such as *Neurod1* (22) (**Suppl. Dataset 8, Figure 5G**). These results indicated that Atoh7 specifically influenced the epigenetic landscape related to the formation of the RGC lineage.

To gain further insight into the mechanisms by which the epigenetic landscape is regulated to achieve RGC-specific gene expression, we performed CUT&Tag for Pou4f2 and Isl1 with E14.5 retinal cells, again in duplicates, and carried out the intersection analysis with the E14.5 enhancers identified by scATAC-seq (**Figure 7A, B**). We identified 1,535 enhancers bound by Pou4f2 (**Suppl. Dataset 6**) and 313 enhancers bound by Isl1 (**Suppl. Dataset 6**). The lower numbers of peaks identified, as compared to Otx2 and Atoh7, likely reflected the more restrictive roles of Pou4f2 and Isl1, rather than experimental variations, as the duplicate experiments were highly reproducible (respective correlation coefficients: 0.853 and 0.867). The two sets of enhancers bound by Pou4f2 and Isl1 respectively were highly RGC specific and most active only in the RGC clusters including early and late RGCs, indicating they indeed mediate the functions of these two RGC specific transcription factors (**Figure 7A, B**). The top enriched motifs as identified by HOMER for the Pou4f2 and Isl1 bound enhancers were consistent with previously reported motifs respectively, again confirming that CUT&Tag identified bona fide binding sites for them (**Figure 7D, E**). These enhancers were also enriched with motifs recognized by other transcription factors, including Dlx1/2, Atoh7, Ebf1-4, and Onecut1/2, indicating that these factors collaborate with Pou4f2 and Isl1. The enrichment of the Atoh7 motif (E box) in the two sets of enhancers indicated many of them were co-bound by Atoh7. Comparison of the three sets indeed identified enhancers co-bound by the three transcription factors in different combinations, including those bound by all three factors (G1), by Atoh7 and Pou4f2 (G2), by Atoh7 and Isl1 (G3), and by Pou4f2 and Isl1 (G4) (**Figure 7C, G**). Of notice is that the activities of the Pou4f2- and Isl1-bound enhancers, irrespective of Atoh7 binding, were markedly reduced, although substantial activities remained as compared to other cell states/types, in the *Atoh7*-null early RGCs (**Figure 7A, B, C, G**). These findings further confirmed that Atoh7 regulated the epigenetic status by influencing the activities of a distinct set of RGC-specific enhancers, likely via both direct and indirect mechanisms, and that other factors function in parallel with Atoh7 to fully activate these enhancers and thereby the associated target genes (22).

**Figure 7.**
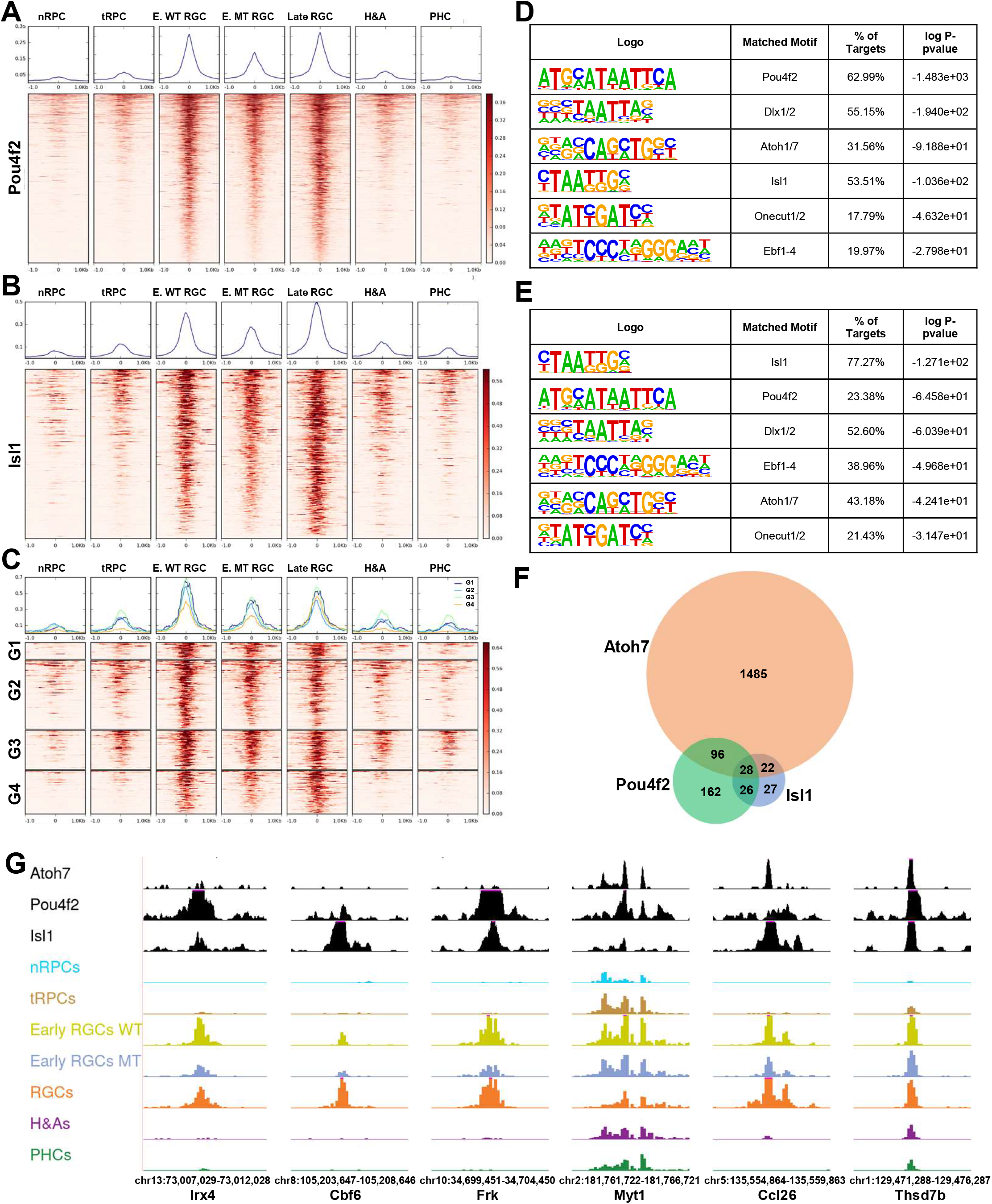
Pou4f2 and Isl1 bind to RGC-specific enhancers. **A**. Activities of Pou4f2-bound enhancers as determined by normalized Tn5 insertion frequencies across different cell states/types at E14.5. **B**. Activities of Isl1-bound enhancers across different cell states/types. **C**. Activities of enhancers co-bound by at least two of the three transcription factors Atoh7, Pou4f2, and Isl1. G1: Atoh7+Pou4f2+Isl1; G2: Atoh7+Pou4f2; G3: Atoh7+Isl1; G4: Pou4f2+Isl1. For **A-C**, genomic regions the MACS2 summits ±1.0 kb were displayed. Color scales represent normalized Tn5 insertion frequencies. Normalized average Tn5 insertion frequencies in the different cell states/types are displayed on top of each heatmap. Pou4f2 summits are used in **C** to generate the heatmap. **D**. Motif enrichment analysis by HOMER of Pou4f2-bound enhancers displayed in **A. E**. Motif enrichment analysis by HOMER of Isl1-bound enhancers displayed in **B. F**. A Venn diagram demonstrating the overlaps of genes likely regulated by Atoh7, Pou4f2, and Isl1. **G**. Example enhancers bound by Pou4f2 and/or Isl1, with or without Atoh7 binding, and their cell state/type-specific activities at E14.5 as displayed in the genome browser. Note that the activities of all these enhancers were highly RGC specific and significantly reduced in the *Atoh7*-null early RGCs (Early RGCs MT) as compared to the wild-type early RGCs (Early RGCs WT). Genome coordinates (mm10) of the displayed regions and the nearby genes are indicated at the bottom.

Many of the genes associated with enhancers bound by Atoh7, Pou4f2, and Isl1 were previously identified downstream genes of these factors, further confirming the validity of our result (**Suppl. Dataset 6**) (22, 37–39). Moreover, these three sets of associated genes overlapped substantially (**Figure 7F, Suppl. Dataset 6**). Since Atoh7, Pou4f2, and Isl1 function in two different but consecutive states of the RGC trajectory, the differential binding of the three transcription factors to individual enhancers and their dynamics along the RGC trajectory allowed for the inference of the different regulatory modes via which individual RGC genes were activated by their sequential and combinatorial actions. These modes included early RGC genes with just Atoh7 binding but little Pou4f2/Isl1 binding such as *Eya2* (**Figure 8, Suppl. Dataset 6**), early RGC genes with both Atoh7 and Pou4f2/Isl1 binding such as *Pou4f2* and *Isl1* (**Figure 9**), late RGC genes with Atoh7 and Pou4f2/Isl1 binding such as *Klf7, Syt13, Ebf3, Pou4f1, Ablim1, Gap43*, and *Elavl4* (**Figure 8, Suppl. Dataset 6**), late RGC genes with Atoh7 binding but without Pou4f2/Isl1 binding such as *Nhlh2, Shh*, and *Ina1*, late RGC genes without Atoh7 binding but with Pou4f2 and/or Isl1 binding such as *Pou6f2* and *Mstn* (**Figure 8, Suppl. Dataset 6**). These modes indicated that Atoh7 is involved in the initial activation of many of the early RGC genes and some late RGC genes. In contrast, Pou4f2 and Isl1 function to sustain the expression of the early RGC genes including themselves, and activate the expression of many late RGC genes, after *Atoh7* is turned off. As indicated by motif enrichment and footprinting analysis (**Figure 4**), other transcription factors, such as Ebf1-4 and Onecut1/2, likely collaborate with Pou4f2 and Isl1 in the process either via shared or distinct enhancers (**Figure 8**) (14, 62, 99). Genes without Pou4f2/Isl1 binding sites were likely just regulated by these other transcription factors; even in these cases, the Atoh7-Pou4f2/Isl1 cascade may still be involved indirectly as genes encoding some of these other transcription factors such as Ebf1-4 and Irx1-6 are subject to its regulation (22, 35, 37, 39). Of notice was that Atoh7 binding sites were also observed in the three SoxC transcription factor genes (**Suppl. Figure 10, Suppl. Dataset 6**), which likely function in parallel with Atoh7 in promoting the RGC lineage from the tRPCs (22, 32, 33). This was particularly the case for *Sox11*, for which multiple Atoh7 and Pou4f2 binding enhancers were identified. These findings indicate that Atoh7 and the SoxC factors likely cross-activate each other in activating the RGC differentiation program.

**Figure 8.**
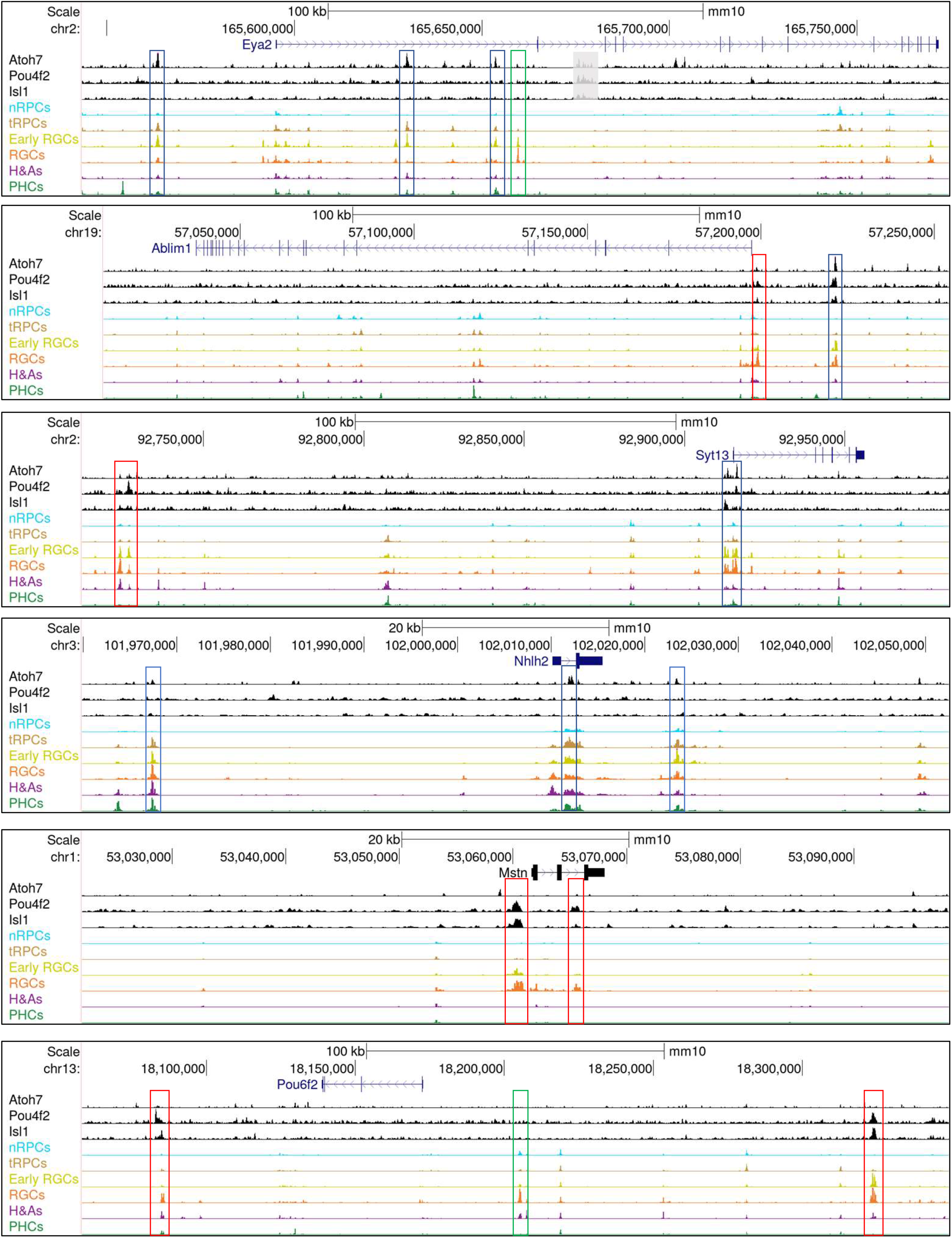
Sequential and combinatorial regulation of RGC-specific genes by Atoh7, Pou4f2, and Isl1. CUT&Tag and E14.5 scATAC-seq genome tracks are displayed for the enhancers associated with six RGC-specific genes, including *Eya2, Ablim1, Syt13, Nhlh2, Mstn*, and *Pou6f2*. All these genes have RGC specific enhancers, but they are bound differentially by the three transcription factors, representing distinct modes of sequential and combinatorial regulation (see text for detail). Enhancers bound by Atoh7 (blue boxes) tend to be activated early (active in tRPCs already) in the RGC trajectory, whereas those just bound by Pou4f2 and/or Isl1 (red boxes) tend to be activated late. RGC-specific enhancers not bound by any of the three transcription factors (green boxes) likely mediate regulation by additional RGC-specific transcription factors.

**Figure 9.**
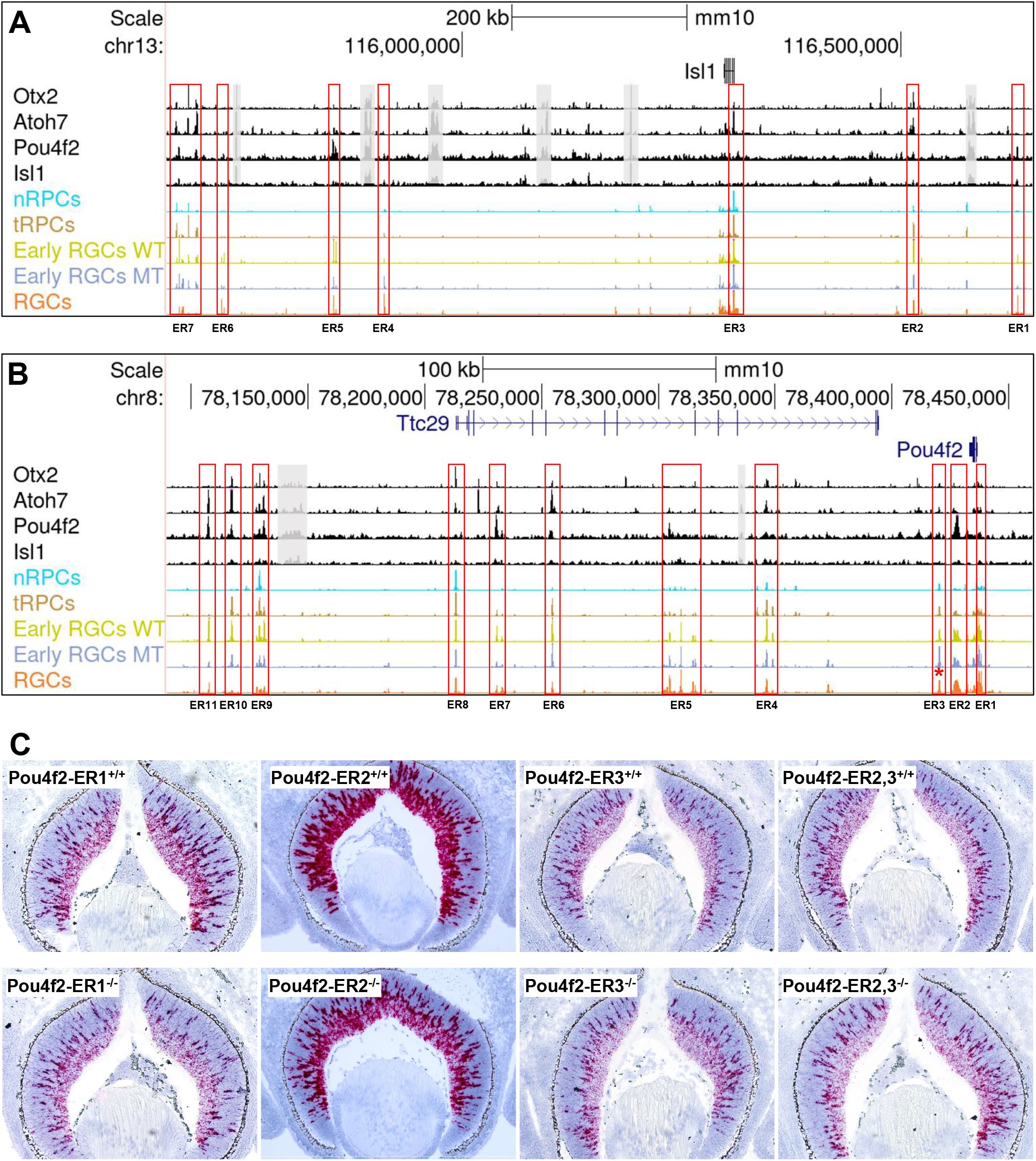
Regulatory mechanisms underlying the early RGC-specific expression of *Isl1* and *Pou4f2*. **A**. Seven putative enhancer regions (ER, red boxes) with distinct dynamics and differential binding by Otx2, Atoh7, Pou4f2, and Isl1 likely regulate the RGC-specific expression of *Isl1*. Displayed are CUT&Tag peaks and E14.5 scATACs-seq peaks surrounding the *Isl1* locus. For clarity, some non-specific peaks falling into repetitive sequence regions and not overlapping with the scATAC-seq peaks were shadowed with gray boxes. **B**. Eleven putative enhancer regions (red boxes) with distinct dynamics and differential binding by Otx2, Atoh7, Pou4f2, and Isl1 likely regulate the RGC-specific expression of *Pou4f2*. Note that most, but not all, *Isl1* and *Pou4f2* enhancers have reduced activities in the *Atoh7*-null early RGCs (blue track) as compared to the wild-type early RGCs (lime track). Enhancer region 3 (ER3, marked by a red asterisk) has increased activity in the *Atoh7*-null early RGCs. **C**. Deletions of three putative *Pou4f2* enhancer regions have different effects on *Pou4f2* expression. *Pou4f2* expression was detected by RNAscope on E14.5 retinal sections with the indicated genotypes. Deletion of ER1 and ER2 leads to reduced *Pou4f2* expression, particularly in the ganglion cell layer, whereas deletion of ER3 or ER2/3 does not change *Pou4f2* expression substantially. *Pou4f2-ER2*^*-/-*^ is a compound heterozygous mouse (*Pou4f2*^*tdtomato/+*^) with both copies of the ER2 elements deleted, whereas *Pou4f2-ER2*^*+/+*^ is a *Pou4f2*^*tdtomato/+*^ control with both ER2 copies intact.

**Figure 10.**
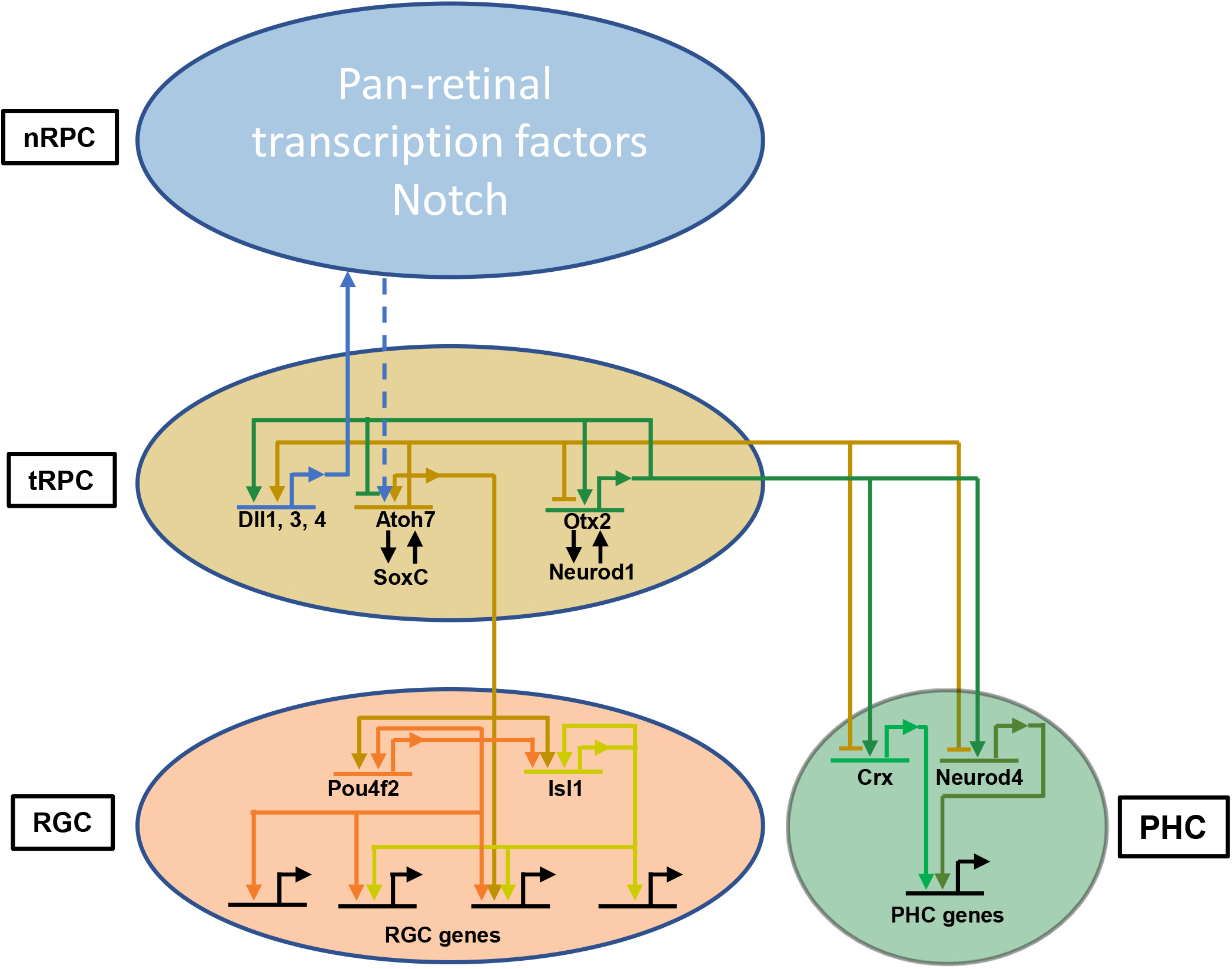
A model on the emergence of the RGC and PHC lineages from transitional RPCs. Establishment of the tRPC state involves the repression of nRPC specific genes, activation of the tRPC-specific transcription factor genes including *Atoh7* and *Otx2*, and downregulation of the Notch pathway. Atoh7 and Otx2 promote expression of Notch ligands, which act on nRPCs to maintain the proliferative and non-differentiating state. In tRPCs, Atoh7 and Otx2, and likely transcription factors for other lineages, repress each other’s expression directly. The cross-repression eventually leads to the dominance of one lineage-specific program in individual cells, which is further stabilized and maintained by downstream lineage specific transcription factors. Isl1 and Pou4f2 are key downstream transcription factors for the RGC lineage, whereas Crx and Neurod1, 4 are key downstream transcription factors for the PHC lineage. RGC-genes receive distinct combinations of inputs from upstream transcription factors in a sequential and combinatorial manner, and thus are subject to different modes of regulation. Similar scenarios likely apply to PHCs and other retinal lineages.

### Epigenetic mechanisms controlling *Isl1* and *Pou4f2* expression

Since *Isl1* and *Pou4f2* are the earliest lineage-specific genes activated during RGC development and their roles in establishing the RGC lineage are well defined (34, 35, 37, 39, 40), deciphering the mechanisms by which these two genes are activated is critical to understanding how RGCs emerge from the multipotent tRPC state. The scATAC-seq and CUT&Tag-data allowed us to examine in detail how these two genes are likely regulated.

Based on the activities of the surrounding enhancers and the locations of the binding sites of Otx2, Atoh7, Pou4f2, and Isl1, seven major putative enhancer regions (ER1-7) likely regulating the expression of *Isl1* were identified in a one million base pair region (**Figure 9A**). The dynamics of the enhancers in these regions across the different cell states/types were in good agreement with the functions of the transcription factors binding them (**Figure 9A**). Thus, the enhancers bound by Atoh7 including ER2, 3, and 7 were all most active in tRPCs and/or early RGCs; some of these enhancers were also bound by Otx2 (ER2) or Otx2 and Pou4f2 (ER7). ER1 and ER5, which were most active in late RGCs, were bound by just Pou4f2 or Pou4f2 and Isl1. Interestingly, ER4 and ER6, which were also most active in late RGCs, were not bound by any of these transcription factors, indicating they likely mediate the function of other transcription factors. The activities of most of these enhancers (ER1, 2, 5, 6, and 7), irrespective of Atoh7 binding, were noticeably reduced in the early *Atoh7*-null (MT) RGCs (**Figure 9A**).

Eleven putative enhancer regions (ER1-11) likely regulating *Pou4f2* were identified, and they were also located over a large genomic region (∼350 kb). Many of these enhancers were located within or beyond the neighboring gene *Ttc29* on the 3’ side of *Pou4f2*. These enhancers likely mediated the expression of *Pou4f2*, instead of *Ttc29*, since their activity dynamics closely mirrored that of *Pou4f2* expression (**Figure 9B**), and *Ttc29* is expressed in the retina at very low levels and not affected in the *Atoh7*-, *Pou4f2*- or *Isl1*-null retinas (22). The binding patterns to these enhancers by the four transcription factors were similar to those observed with the *Isl1* enhancers, and the dynamics of these enhancers was also closely related to the functions of the transcription factors binding them. Thus, the *Pou4f2* enhancers included those bound by Atoh7 (ER4, 6, 8, 9, 10, 11), with or without Otx2 and/or Pou4f2 binding, those with just Pou4f2 binding but little Atoh7 binding (ER2, 5 and 7), and those not bound by any of these transcription factors (ER1, 3). No substantial binding for Isl1 was observed in these enhancers. Like the *Isl1* enhancers, the *Pou4f2* enhancers bound by Atoh7 (ER4, 6, 9, 10, 11) were most active in tRPCs and/or early RGCs, whereas those bound by just Pou4f2 (ER2, 5, and 7) were most active in early and/or late RGCs. Again, the activities of many of these enhancers including ER2, 4, 7, 8, 9, 10, and 11, irrespective of Atoh7 binding, were markedly reduced in the early *Atoh7*-null RGCs (**Figure 9B**).

The features of the enhancers associated with *Isl1* and *Pou4f2* suggested that the two genes are activated through similar sequential functions of Atoh7 and Pou4f2/ Isl1, with Atoh7 acting first by competing with Otx2 and binding to select enhancers associated with the two genes, leading to their initial activation. Once expressed, Pou4f2, Isl1, and likely other transcription factors then bind to distinct but overlapping sets of enhancers to maintain their expression. The contributions of the individual candidate enhancers to *Pou4f2* and *Isl1* expression could be tested experimentally. As a first step, we deleted three *Pou4f2* enhancer regions (ER1-3) by CRISPR/cas9 and examined the effects on *Pou4f2* expression by RNA-scope in situ hybridization at E12.5, E14.5, and E17.5 (**Figure 9B, C, Suppl. Figure 11**). ER1 was in the promoter region and upstream of a presumptive TATA box, not bound by any of the four transcription factors, and most active in RGCs (**Figure 9B**). Homozygous deletion of ER1 (2,543 bp) markedly reduced *Pou4f2* expression at E14.5 and $17.5, but not E12.5 (**Figure 9C**). ER2 was located downstream of the gene body, extensively bound by Pou4f2, and most active in RGCs (**Figure 9B**). Mice with both alleles of ER2 deleted exhibited substantially reduced *Pou4f2* expression in the retina at E14.5 (**Figure 9C**), but not E12.5 or E17.5 (**Suppl. Figure 11**). In both ER1 and ER2 deletions, *Pou4f2* expression still occurred with the normal spatial and temporal patterns, but the reduction was most prominent in the future ganglion cell layer (**Figure 9C**), indicating that the other enhancers participate in its regulation. ER1 and ER2 may not be required for the initial activation, but the maintenance, of *Pou4f2* expression, and in the ER2 deletion, the function of ER2 may be compensated by other enhancers at later stages. ER3 was located further downstream, active in all cell states/types, and not bound by any of the four transcription factors. Deletion of ER3 (1,419 bp) did not lead to appreciable changes in *Pou4f2* expression at any stage (**Figure 9C, Suppl. Figure 11**). Interestingly, when the continuous region encompassing both ER2 and ER3 (10,017 bp) were deleted, no obvious changes were observed in *Pou4f2* expression either (**Figure 9C, Suppl. Figure 11**). The differences between ER2 deletion and ER2/3 deletion indicated that ER3 likely served as a repressor. This was consistent with the observation that the activity of ER3 was increased in the *Atoh7*-null retina (**Figure 9B**). These results indicated that the final gene expression output, namely the spatial and temporal expression of *Pou4f2* in RGCs, resulted from the integrated and likely redundant inputs of multiple enhancers.

## DISCUSSION

In this study, we use scATAC-seq and CUT&Tag to investigate the shifting epigenetic landscape in the early developing retina at single cell resolution and to gain insights into the mechanisms by which key transcription factors interact with and influence the epigenetic landscape and promote retinal cell differentiation toward individual lineages, particularly RGCs. The scATAC-seq data allowed us to group retinal cells from two stages (E14.5 and E17.5) based on the differential accessibility/activity of enhancers across different cell states/types. The clustering results are highly consistent with those obtained with scRNA-seq and reveal the same developmental trajectories of the different lineages in early retinal development (8, 21, 22), indicating that the gene expression changes along the developmental trajectories of individual lineages are associated with, and likely resulted from, shifts in the epigenetic landscape. Accordingly, the differentially active enhancers associated with the distinct cell states allow for probing the regulatory mechanisms controlling retinal cell fate specification and differentiation.

Along individual trajectories from nRPCs to tRPCs, and then to fate-specified neurons, cell state/type-specific enhancers become activated or inactivated, resulting in corresponding changes in gene expression. Thus, putative enhancer elements regulating cell state/type specific expression of individual genes can be identified based on the correlation of the gene expression and enhancer activity dynamics. The epigenetic landscape and thereby the correlating gene expression are determined by transcription factors expressed in individual cell states/types. Supporting this, we found that DNA motifs for many transcription factors known to function in the developing retina are enriched, and corresponding footprints are present, in enhancers active in distinct but often consecutive cell states along individual trajectories. Transcription factors functioning at one state likely also drive the cells to the next state, as many enhancers often are activated already at low levels in the state before the gene is expressed. Our findings are consistent with a couple of recent reports (69, 100) and lay the foundation for further investigating how key transcription factors interact with the epigenome to regulate retinal cell differentiation. To that end, we examined the changes in the epigenetic landscape in the absence of Atoh7 and used CUT&Tag to identify enhancers bound by four cell state/type-specific transcription factors, Otx2, Atoh7, Pou4f2, and Isl1, which revealed their sequential and combinatorial action in shaping the epigenetic landscape toward distinct cell fates, particularly the RGC fate. Although our CUT&Tag data are not at single cell resolution due to the current limitations of the technology, intersection of CUT&Tag with scATAC-seq data revealed the dynamics of the enhancers mediating the function of these four transcription factors, the overlaps of the distinct sets of enhancers, and the genes these enhancers are associated with, which strongly support a general paradigm by which key transcription factors both collaborate and compete to drive tRPCs toward distinct lineages **(Figure 10**).

Particularly, insightful observations were obtained for the critical multipotent tRPC state which is poised to commit to one of the retinal cell fates (22). The most prominent epigenetic characteristic of this state is the activation of enhancers containing E box motifs, which are recognized by bHLH transcription factors including Atoh7. As suggested by extensive previous studies (101–109), interactions between the Notch pathway and the bHLH transcription factors likely are the major underlying mechanism for this epigenetic change; downregulation of the Notch pathway by lateral inhibition is critical for activation of the bHLH genes and retinal cell differentiation, whereas bHLH transcription factors activate the Notch ligand genes, which in turn activate the pathway in nRPCs to promote proliferation and inhibit differentiation (94, 95, 100, 102, 110, 111). Our finding that Atoh7 regulates multiple component genes of the Notch pathway fits this model. Multiple bHLH transcription factors are involved in distinct retinal cell fates, Atoh7 and Neurog2 in RGCs, Neurod1 and Neurod4 in PHCs, and Bhlhe22 and Ptf1a in H&As (18, 84–90). The distinct functions of these bHLH factors are also manifested by the cross repression between some of these genes. For example, Atoh7 represses *Neurod1* and *Neurog2*, whereas Ptf1a represses *Atoh7* (22, 87, 112). Many of these factors are co-expressed in tRPCs, and some of them continue to be expressed in the differentiated neurons. Although these bHLH factors belong to distinct classes and have some sequence preferences for the E box motif, their binding specificities are significantly redundant (113–116). Thus, the molecular basis for the cell lineage specificities of individual bHLH factors is not known. Elucidation of these specificities will require discovery and comparison of the in vivo binding sites of multiple bHLH factors, biochemical dissection of their intrinsic properties, and identification of their interacting partners.

Other than the bHLH factors, multiple other transcription factors involved in distinct cell fates, including Otx2, Foxn4, Sox4/Sox11, and Onecut1/2, are co-expressed in tRPCs (22). Our findings and previous reports indicate that many of these factors such as Otx2, Foxn4, and Sox4/11 also interact with the Notch pathway (33, 111). Consistent with the two factors being required in two distinct lineages, Atoh7 in RGCs and Otx2 in PHCs, their target genes included those respectively involved in the differentiation, structure, and physiological function of the two cell types, including *Atoh7* and *Otx2* themselves. Importantly, Atoh7 and Otx2 also bind mutually to each other’s enhancers and enhancers associated with genes of the opposite fate. These findings indicate that Atoh7 and Otx2 also cross-regulate genes of the opposite lineage directly (**Figure 10**). Given the cell type specific roles of Atoh7 and Otx2, the cross-lineage regulation is likely repressive. Thus, Atoh7 and Otx2 likely both collaborate and compete in regulating downstream genes, depending on the gene contexts; they collaborate in regulating genes with general functions in multiple lineages, but compete in regulating lineage-specific genes by promoting genes for one lineage and repressing genes for the other. The competition between Otx2 and Atoh7 is best exemplified by their co-regulation of the PHC-specific gene *Neurod1*, Otx2 being activating whereas Atoh7 being repressive, via shared enhancers. The target genes of Atoh7 and Otx2 also included those involved in the H&A lineage. We previously proposed that transcription factors for different lineages collectively determine the temporal competence of tRPCs and compete in this shared cell state to drive them to distinct trajectories (22). This idea of competition between transcription factors for different lineages was further elaborated recently by integration of scRNA-seq and scATAC-seq data and network modeling (69). Our current findings provide direct experimental evidence to support this general paradigm; the collaboration and competition between transcription factors promoting different lineages lead to dominance of individual gene expression programs and emergence of definitive cell lineages. This is further supported by previous findings that deletions of genes encoding these factors often lead to the increase of other fates; deletion of *Atoh7* causes increased photoreceptors and amacrine cells (29, 30), deletion of *Foxn4* leads to increased production of RGCs and photoreceptors (111, 117), whereas deletion of *Otx2* results in fate switch to amacrine and RGC fates (70, 118). However, how the interaction among these lineage-specific transcription factors in tRPCs leads to the eventual dominance of just one unambiguous genetic pathway and thereby commitment to the corresponding cell fate remains to be elucidated.

Nevertheless, our study already offers insights into how RGC-specific gene expression is achieved and the RGC fate is specified. Comparison of the wild-type and *Atoh7* mutant retinal cells suggested that Atoh7 is not required for establishing the overall tRPC epigenetic landscape. This may be due to the presence of other transcription factors expressed in tRPCs, which may collectively establish this state. For those enhancers normally bound by Atoh7 but still fully accessible (active) in the absence of Atoh7, Atoh7 may be just one of multiple transcription factors interacting with them. On the other hand, Atoh7 is required for the full activities of many enhancers associated with genes critical for RGC differentiation and function. Atoh7 likely exerts its effects on the epigenetic landscape both directly and indirectly, as not all Atoh7-dependent enhancers are bound by Atoh7. Enhancers dependent on Atoh7 but not bound by it likely are activated by downstream transcription factors such as Pou4f2 and Isl1. The activity dynamics of these enhancers along the RGC lineage trajectory is consistent with the differential binding of Atoh7 and Pou4f2/Isl1 to them as well as their stage-specific functions; Atoh7 tends to bind to enhancers active in tRPCs and early RGCs (RGC precursors) and thus are involved in the initial activation of the associated genes when the RGC fate is specified, whereas Pou4f2 and Isl1 tend to bind to enhancers most active in early and late RGCs and likely are involved in activating and maintaining genes expressed in fate-specified RGCs. Thus, Atoh7 and key downstream transcription factors (14), including Pou4f2, Isl1, and likely the Ebf factors (Ebf1-4), the Irx factors (Irx1-6), and Onecut1/2, regulate RGC-specific genes in a sequential and combinatorial fashion. Individual RGC genes receive distinct combinations of regulatory inputs; some require both Atoh7 and the downstream transcription factors, whereas others require only the downstream factors (**Figure 10**). We previously proposed that the SoxC factors may also serve as key upstream positive regulators in tRPCs based on their expression patterns, their roles in RGC development, and the findings that the RGC lineage still forms in the absence of Atoh7 (22, 32, 33). Our findings that the SoxC DNA motifs are highly enriched in the Atoh7 bound enhancers, and that many RGC-specific enhancers can still be activated at substantial although much reduced levels in the *Atoh7*-null retina in an RGC-specific fashion, further support this idea.

As the earliest genes specifically activated in the RGC lineage, *Pour4f2* and *Isl1* activation coincides with the emergence of the RGC lineage. Pou4f2 and Isl1 are also critical for activating and maintaining the RGC-specific gene expression program and stabilizing the lineage (34, 37, 39). Thus, deciphering how *Pour4f2* and *Isl1* are regulated is critical for understanding the establishment of RGC fate. We have identified multiple putative enhancers for each gene with distinct dynamics of activity and transcription factor binding profiles, which allow us to infer the regulatory mechanisms by which they are activated and maintained. Consistent with the two genes overlapping completely in developing RGCs, their regulation seems to follow very similar scenarios. The binding of Atoh7 to select enhancers in *Pou4f2* and *Isl1* likely is the first step toward the RGC fate. Atoh7 must overcome the repression of Otx2, and likely other non-RGC lineage-specific factors, to activate *Pou4f2* and *Isl1* by directly repressing *Otx2* and these other factors and by competing with them directly at select *Pou4f2* and *Isl1* enhancers (**Figure 10**). As discussed above, the SoxC factors may also be involved in their initial activation. Once expressed, Pou4f2 and Isl1 bind to select enhancers of the two genes, supporting previous conclusions that the Pou4f2 and Isl1 proteins maintain their own expression through auto- and cross-regulation once *Atoh7* is turned off (34). The maintenance of *Isl1* and *Pou4f2* expression likely also requires other transcription factors, since some RGC-specific enhancers are not bound by either Pou4f2 or Isl1. Of note is that for both *Pou4f2* and *Isl1*, their enhancers scatter over large genomic regions, and individual enhancers likely only contribute to specific aspects of the final expression outcome. Similar findings have been reported for enhancers regulating motor neuron-specific genes (119). Experimental validation of the function of these enhancers, as we have begun to do with *Pou4f2*, will shed light on how these two genes are activated and how the RGC fate is specified, although redundancies among different enhancers may pose a challenge. Future efforts should focus on understanding how inputs from multiple transcription factors are integrated via these enhancers both quantitatively and quantitatively to achieve the eventual expression outputs.

## ACCESSION NUMBERS

The E14.5 and E17.5 scRNA-seq data, the E14.5 and E17.5 saATAC-seq data, and the CUT&Tag data have all been deposited into the NCBI Gene Expression Omnibus with an accession number GSE220587. The data are available to the public without restrictions.

## Supporting information

Supplementary figures, figure legends and other info

## SUPPLEMENTARY DATA

## ACKNOWLEDGEMENT

We thank other members of the Mu laboratory and members of the Department of Ophthalmology, University at Buffalo, for helpful discussions. We thank the Flow and Image Cytometry Shared Resource at the Roswell Park Comprehensive Cancer Center for cell sorting. CRISPR-mediated deletions of the *Pou4f2* enhancers were generated with the help of Aimee Stablewski and Dawn Barnas at the Gene Targeting and Transgenic Resource at the Roswell Park Comprehensive Cancer Center. Construction and sequencing of scATAC-seq and scRNA-seq libraries were carried out at the Genomics and Bioinformatics Core of the University at Buffalo. Bioinformatics analysis was carried out using facilities at the Center for Computational Research at the University at Buffalo.

