## Supplementary figures, figure legends and other info for "Key transcription factors influence the epigenetic landscape to regulate retinal cell differentiation"

#### Supplementary Figure Legends

**Suppl. Figure 1.** Scatter plots of FACS for the isolation of six different cell populations used in scATAC-seq. Note the Low gating threshold we used which allow these populations to contain cells in different but continuous cell states along individual lineage trajectories. See text for details.

**Suppl. Figure 2.** Quality evaluation of scATAC-seq data from the seven purified cell samples from two developmental stages. Top row are E14.5 samples and bottom row are E17.5 samples. Rates of enrichment of scATAC-seq fragments around transcription start sites (left), distributions of scATAC-seq fragment sizes (middle), and their localizations around transcription start sites (center) were all in good agreement with previously published data.

**Suppl. Figure 3.** Feature heat map indicating the gene activities as determined by GeneScore of example marker genes in distinct E14.5 clusters. *Ccnd1*, nRPCs; *Gadd45a*, tRPCs; *Pou4f2*, early and late RGCs; *Sncg*, late RGCs; *Ptf1a*, H&As; *Crx*, PHCs.

**Suppl. Figure 4.** Feature heat map gene activities as determined by GeneScore in distinct E17.5 clusters. *Fgf15*, nRPCs; *Gadd45a*, tRPCs; *Pou4f2*, RGCs; *Sncg*, RGCs; *Ptf1a*, H&As and SACs; *Crx*, PHCs; *Isl1*, RGCs and SACs. The lower right panel shows the contribution of different FACS purified cell samples to the different clusters. Although not as specific as the E14.5 data, the *Atoh7*-null sample (MT zsGreen) did not contribute to the RGC cluster.

**Suppl. Figure 5.** scRNA-seq with the same E14.5 cell populations generate the same clusters as scATAC-seq. UMAP projections of E14.5 scATAC-seq (left) and scRNA-seq (right) are placed side by side, and corresponding clusters are marked and color-coded the same way.

**Suppl. Figure 6.** UMAP clustering of all the wild-type cells from E14.5 and E17.5. Note that the clustering structure is very similar to when E14.5 and E17.5 cells are clustered separately (compare with Figure 1). Cluster identities are determined by GeneScore activities of marker genes (not shown). Corresponding clusters from E14.5 and E17.5 are separated, but always positioned right next to each other. SACs (starburst amacrine cells are only present in E17.5 cells).

**Suppl.** Figure 7. Putative enhancers in example genes as identified by P2G analysis. **A.** *Nefm* and *Nefl* are likely co-regulated by multiple enhancers as indicated by the P2G linkages (as indicated by the arched lines linking the enhancers to the transcription start site). These enhancers all have the strongest activities in RGCs, the cell type in which *Nefm* and *Nefl* are expressed. **B.** Distinct enhancers regulating the cell type-specific expression of Onecut1 are identified, including those regulating its expression in tRPCs (tan), RGCs (orange), PHCs (green), and H&As (purple). **C-E.** Three genes involved in proliferation (*Pcna*, *Mcm2*, *Mcm6*) have no associated enhancers whose activities mirror their cell state-specific expression in RPCs. Although multiple putative enhancers are present in the vicinity of these genes, these enhancers are open in all cell states and display no cell state/type specificity related to their expression.

**Suppl. Figure 8.** tRPC-specific regulation of genes involved in distinct lineages. Example genes regulating distinct retinal lineages, including *Foxn4* or H&As, *Neurog2* and *Sox11* for RGCs, *Olig2* for cones and horizontal cells all possess putative tRPC specific enhancers (red boxes, not all are marked). *Dll1*, which encodes a Notch ligand, also has tRPC-specific enhancers, reflecting its expression in this cell state.

**Suppl. Figure 9.** Peak to gene analysis of differentially accessible peaks between wild-type (WT) and *Atoh7*-null (MT) cells in the early RGC clusters. To perform the analysis, WT and MT cells in the scATAC-seq and scRNA-seq data were separated and compared. Note the high correlation between the two sets of data, indicating the changes in the epigenetic status underlie those in gene expression.

**Suppl. Figure 10.** Atoh7 regulates the SoxC genes. This is particularly the case for *Sox11*, which is associated with multiple enhancers strongly bound by Atoh7 (red boxes). *Sox4* and *Sox12* likely are also regulated by Atoh7 but to lesser degrees.

**Suppl. Figure 11.** RNAscope in situ hybridization of Pou4f2 at E12.5 and E17.5 of different enhancer deletion mutants. These are the same mutant as shown in Figure 9C. Control and mutant pairs are from littermate embryos. There are some slight stage differences of the E12.5 embryos among the different genotypes.

### Suppl. Figure 1

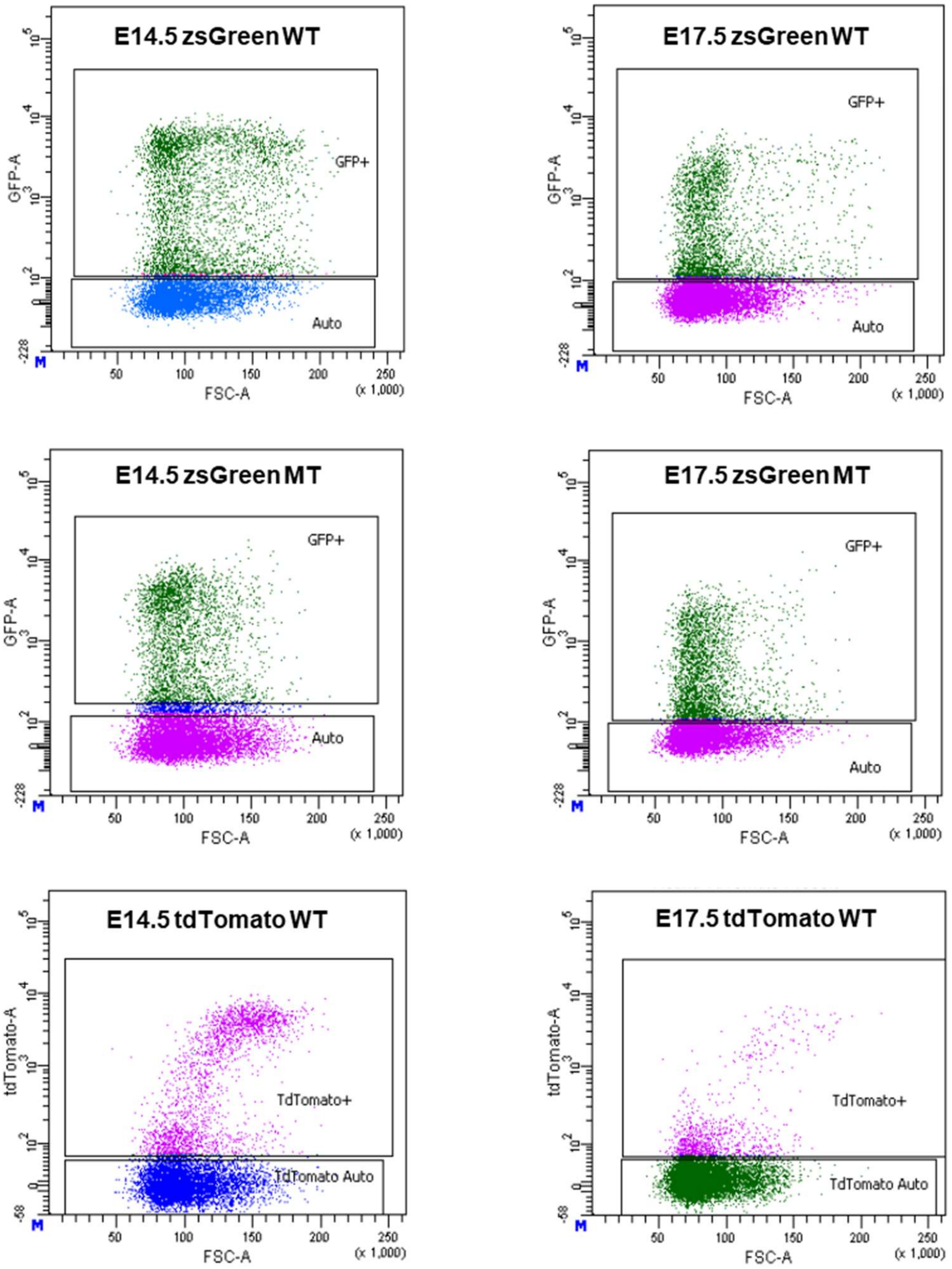

### Suppl. Figure 2

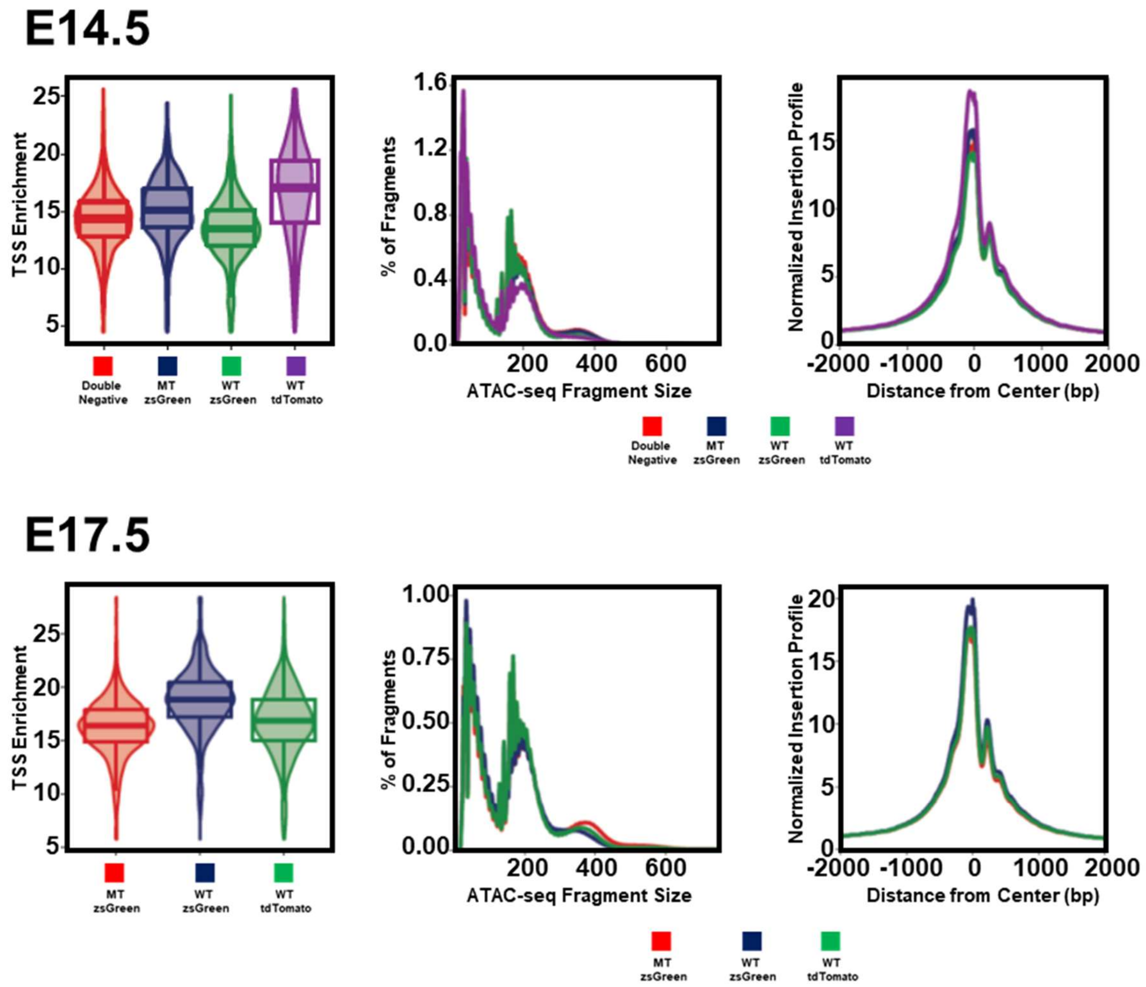

### Suppl. Figure 3

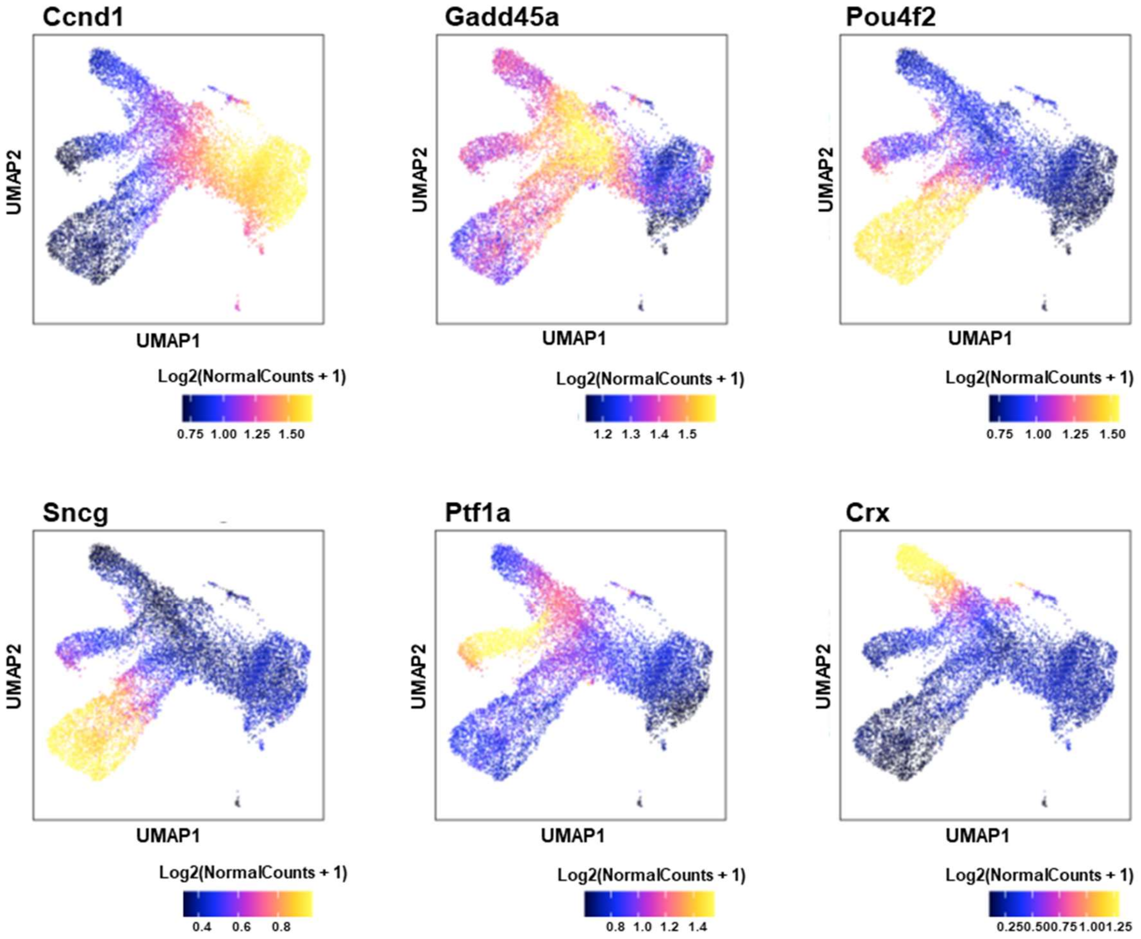

### Suppl. Figure 4

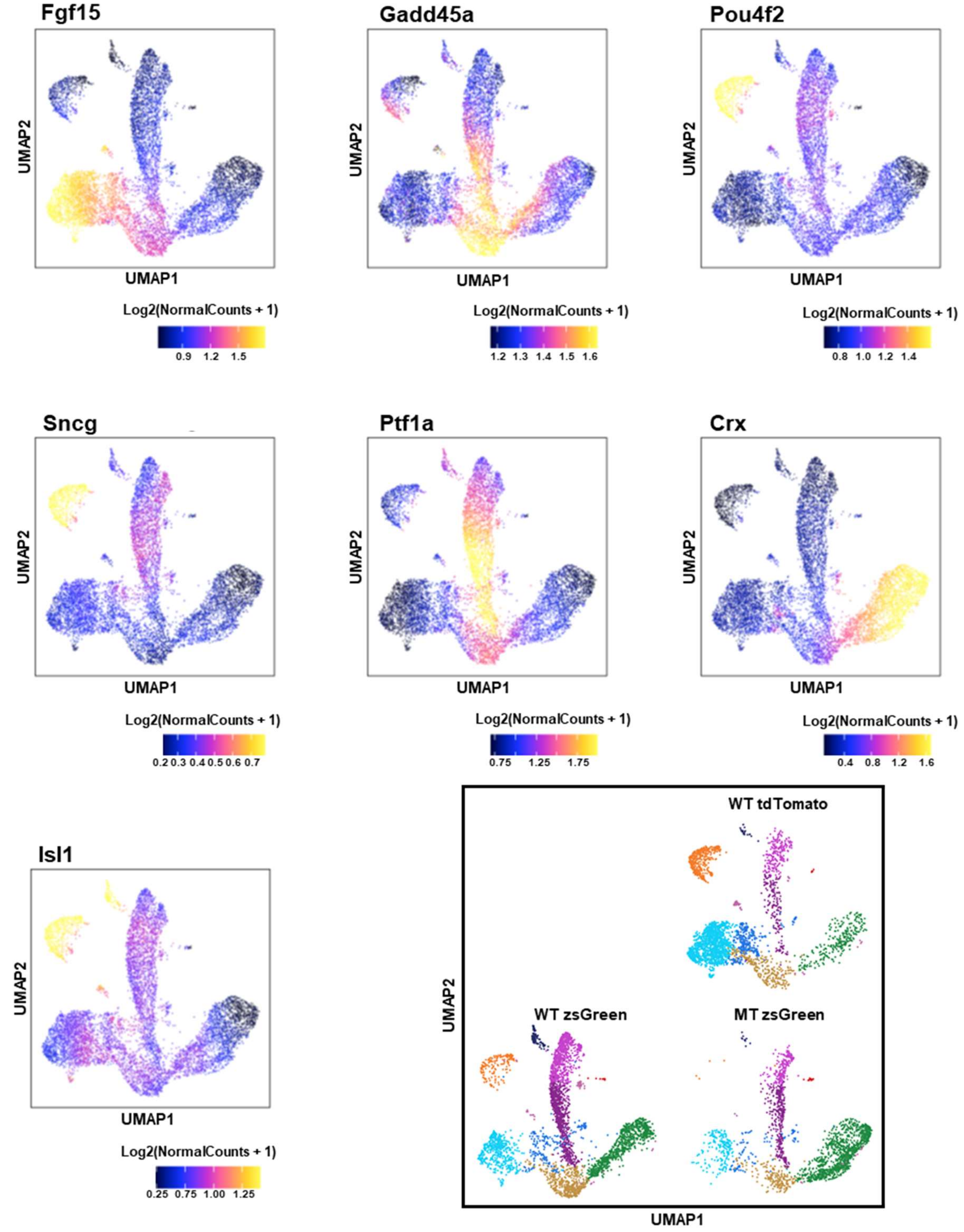

### Suppl. Figure 5

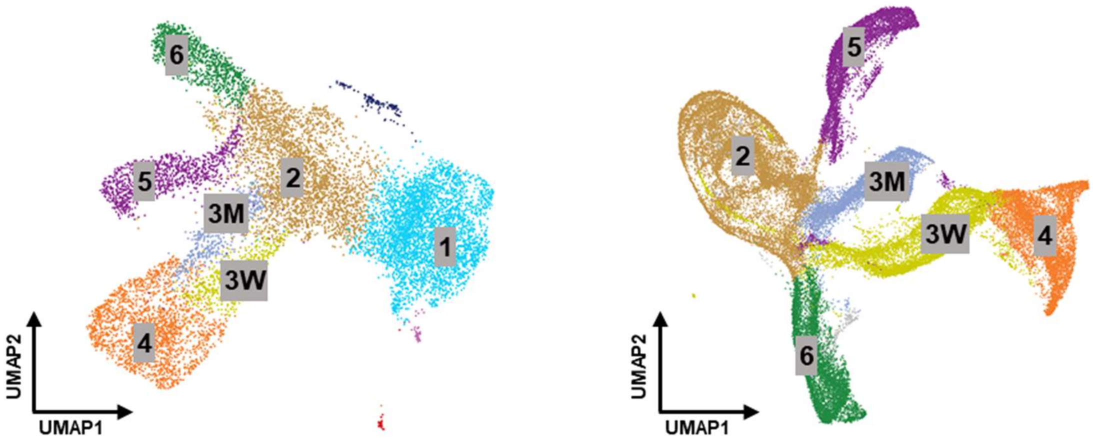


1: Naive RPCs; 2: Transitional RPCs; 3W: Early RGCs (WT);
3M: Early RGCs (MT); 4: RGCs; 5: H&A; 6: PHCs

### Suppl. Figure 6

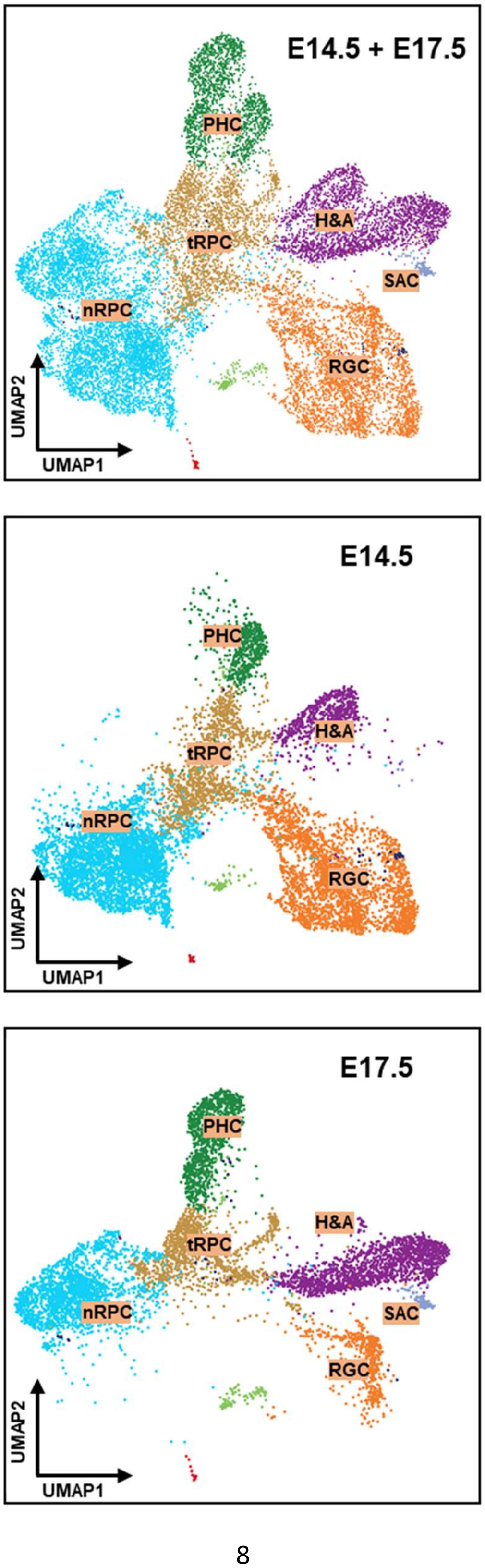

#### Suppl. Figure 7

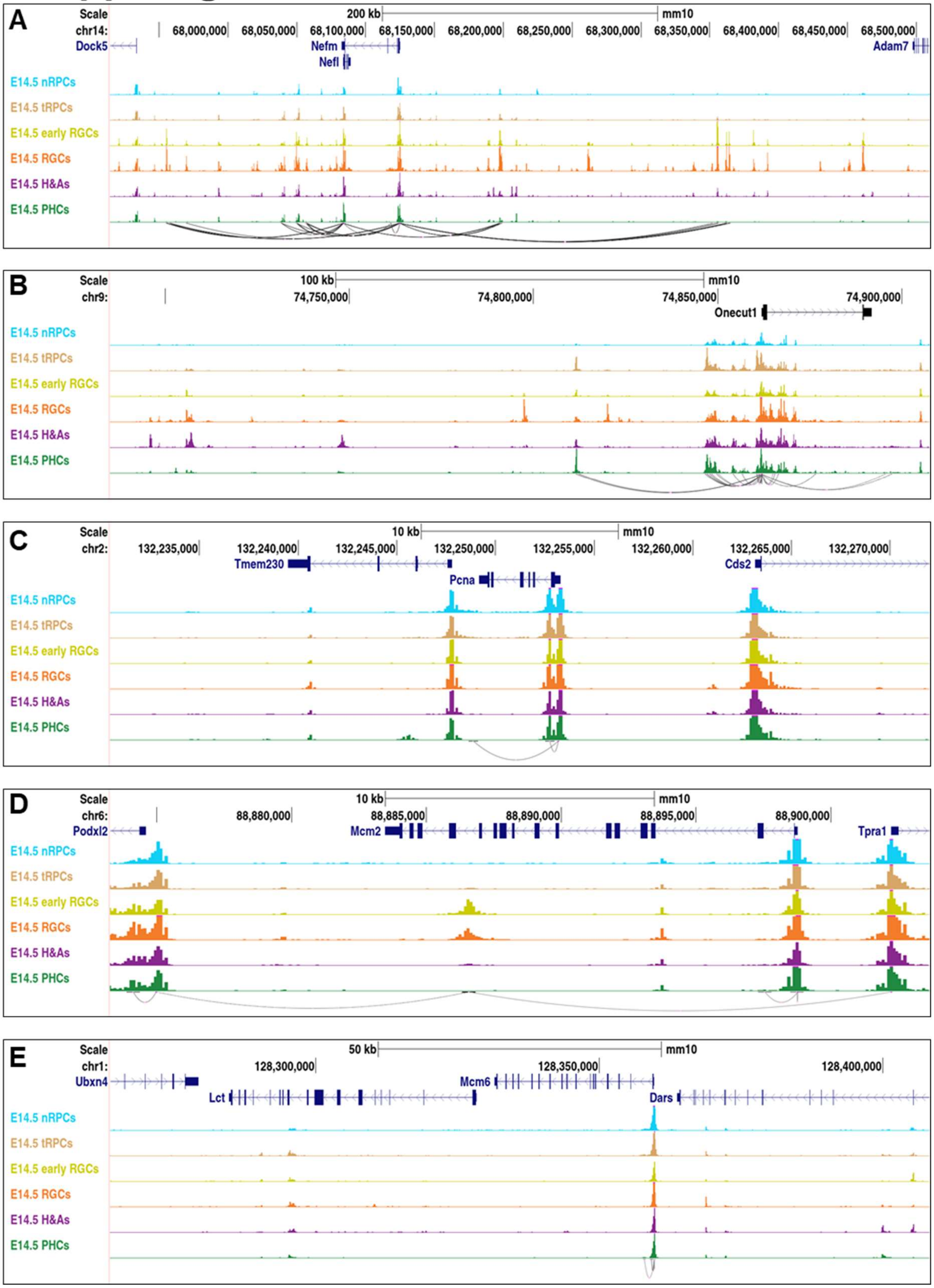

### Suppl. Figure 8

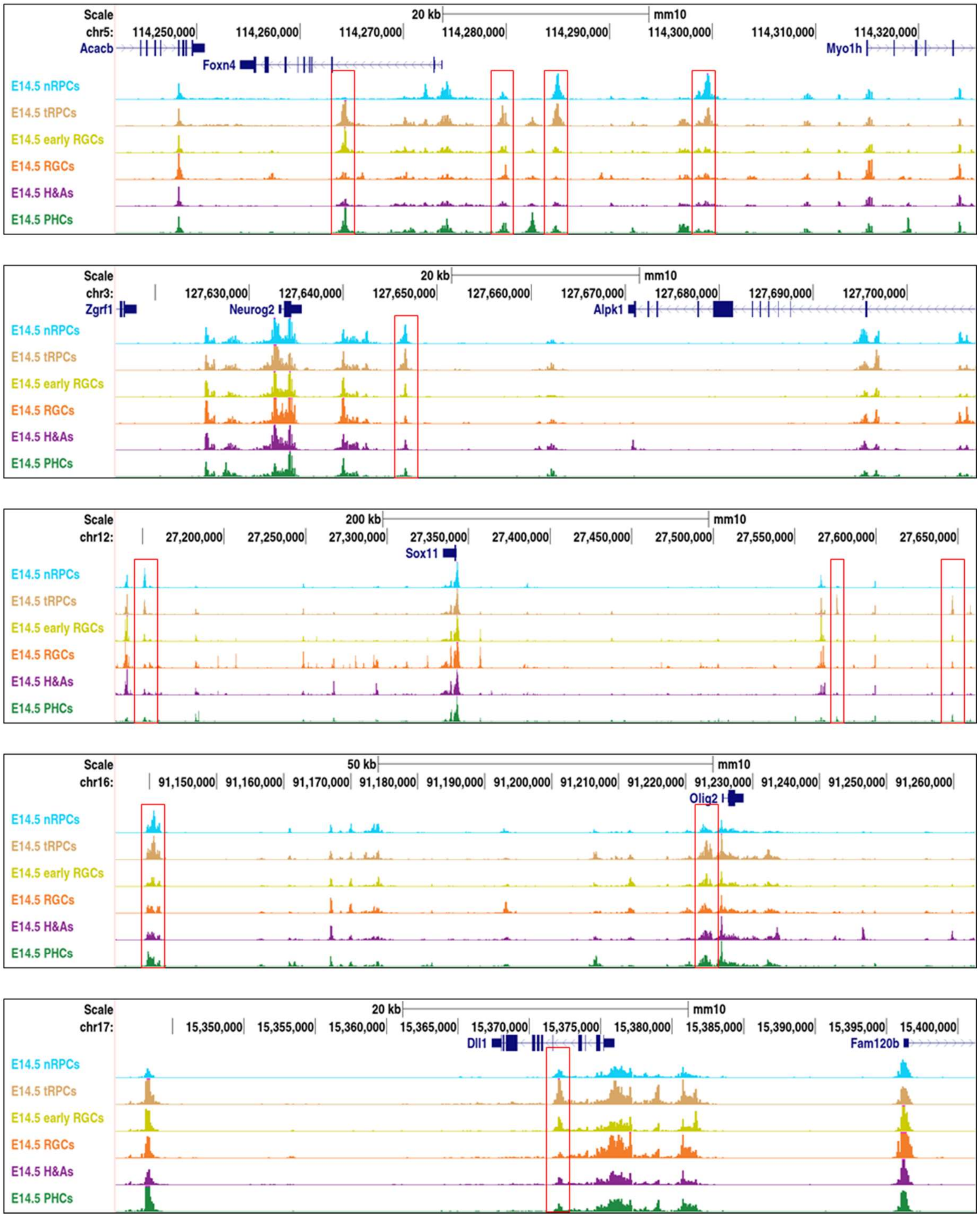

#### Suppl. Figure 9

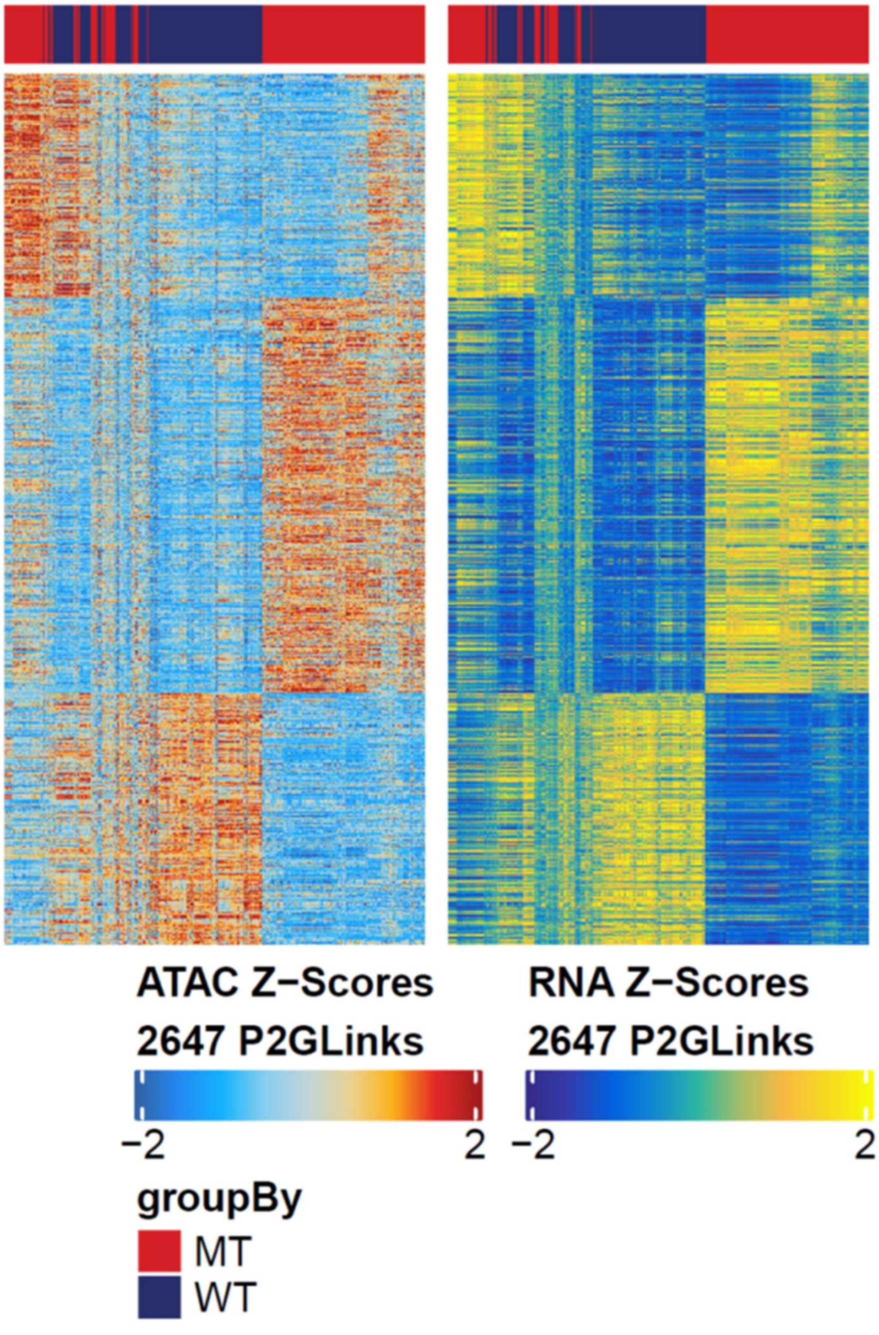

### Suppl. Figure 10

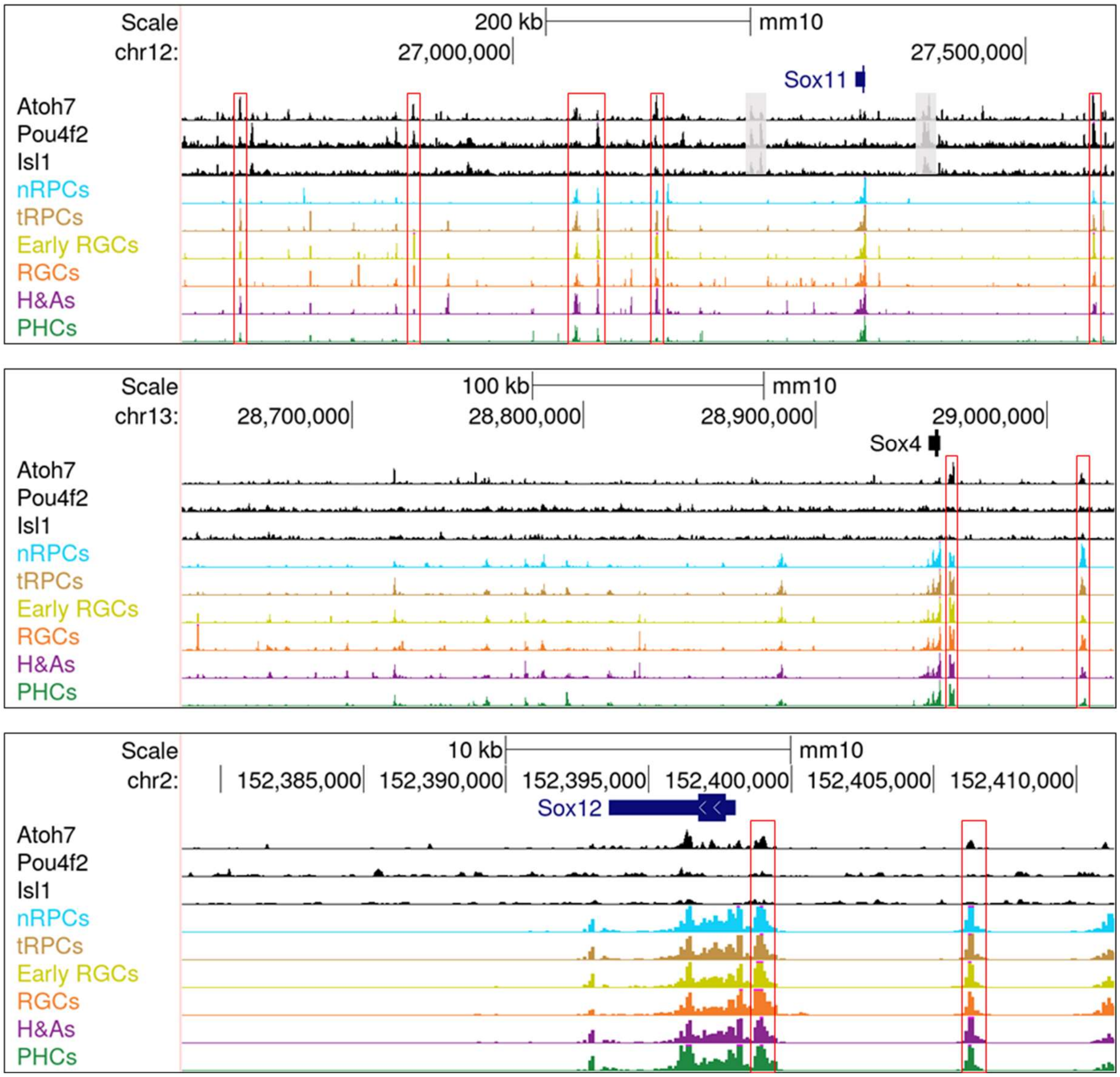

### Suppl. Figure 11

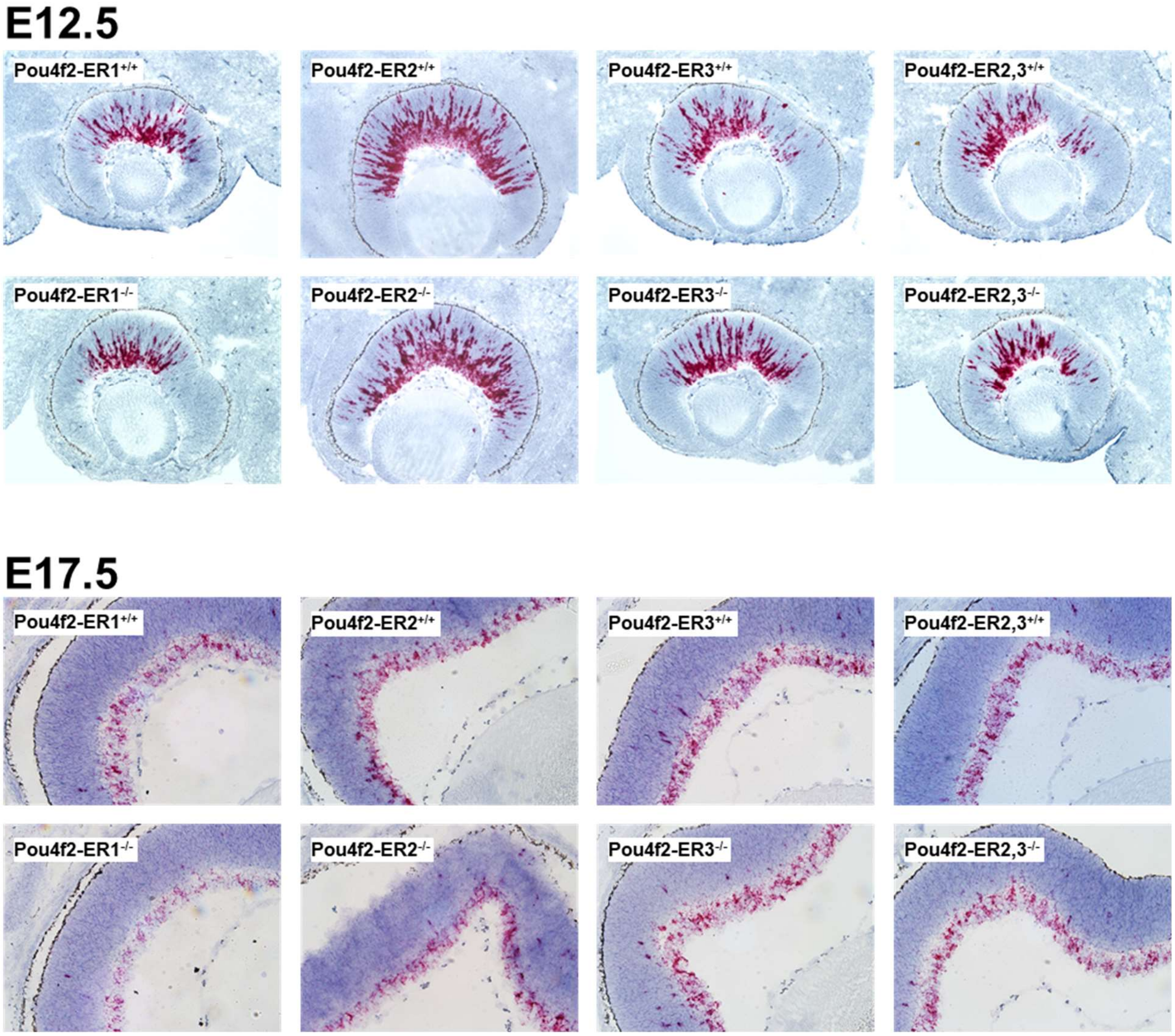

##### Cell numbers in individual scATAC-seq samples

| sample | Estimated number of cells | Median fragments per cell | Fraction of fragments overlapping any targeted region | Fraction of transposition events in peaks in cell barcodes |
|---|---|---|---|---|
| **Atoh7-zsGreen-Lacz-E14_5** | 3782 | 26648 | 71.50% | 56.30% |
| **Atoh7-zsGreen-wt-E14_5** | 2817 | 62645 | 66.30% | 51.00% |
| **Pou4f2-tdTomato-E14_5** | 2789 | 30720 | 71.10% | 58.50% |
| **Atoh7_Pou4f2-double_negative-E14_5** | 3514 | 31764 | 69.50% | 52.00% |
| **Atoh7-zsGreen-Lacz-E17_5** | 1879 | 46681 | 73.30% | 60.30% |
| **Atoh7-zsGreen-wt-E17_5** | 3492 | 26508 | 78.20% | 70.30% |
| **Pou4f2-tdTomato-E17_5** | 2745 | 32756 | 75.60% | 63.00% |

**Genome coordinates for Figures 2E**

Sox2
chr3:34,441,049-34,446,048

Vsx2
chr12:84,530,188-84,535,188
chr12:84,539,211-84,544,211

Zfp36l1
chr12:80,060,453-80,065,453

Fgf15
chr7:145,007,607-145,012,607

Sfrp2
chr3:83,580,762-83,585,762

Atoh7
chr10:63,087,488-63,092,488
chr10:63,096,419-63,101,419

Neurod1
chr2:79,377,021-79,382,021
chr2:79,397,256-79,402,256

Otx2
chr14:48,578,466-48,583466
chr14:48,737,454-48,742,454

Pou4f2
chr8:78,425,481-78,430481

Pou4f1
chr14:104,469,437-104,474,437

Gap43
chr16:42,253,848-42,258,848
chr16:42,267,084-42,272,084

Rbpms
chr8:33,775,949-33,780949

Pou6f2
chr13:18,321,830-18,326829

Ptf1a

chr2:19,429,093-19,434093

Tfap2a
chr13:40,702,176-40,707,176

Crx
chr7:15,862,327-15,867,327
chr7:15,904,879-15,909,879

Neurod4
chr10:130,273,580-130,278,580
chr10:130,285,932-130,290,932

#### crRNA target sequences for *Pou4f2* enhancer deletions

The following are target sequences in the crRNAs used to delete the *Pou4f2* enhancer regions. Four crRNAs were used to delete each enhancer region by CRISPR. To delete ER2 and 3, Enh351, Enh332, Enh231, and Enh232 were used.

**ER1**

```
>enh151
ATTTAGGTGACTGGCACTGC

>enh131
AATTCCCCCAGACTGTTACA

>enh152
TAAGGTGCACCTCACAGTTC

>enh132
ACCAAGGAGGGAAACCGCAT
```

**ER2**

```
>enh251
ATCCAAGCCTGCAAAAGATT

>enh231
CCACCTAGCCCTACATTTGG
>enh252

ATATTCCCGTGTACAAGGAT

>enh232
GACCTGCCTAGAAAGTTGGA
```

**ER3**

```
>enh351
TTGAGGACCTGTTCATCACG

>enh331
ATCAGTGAATCTTGCTACCA

>enh332
TTCATCACGAGGCTTTAATC

>enh352
GGGAGATGTAACTAAACCCC
```

**Genome coordinates for *Pou4f2* enhancer deletions**

Pou4f2-ER1: chr8:78,436,978-78,439,520 (2,543 bp)

Pou4f2-ER2: chr8:78,425,525-78,429,770 (4,246 bp)

Pou4f2-ER3: chr8:78,419,739-78,421,157 (1,419 bp)

Pou4f2-ER2,3: chr8:78,419,755-78,429,771 (10,017 bp)

#### Genotyping primers for the *Pou4f2* enhancer deletion alleles

| Enhancer Regions | Name | Sequence | Name | Sequence | Product Length (bp) |
|---|---|---|---|---|---|
| ER1 | Enh1-5up | TGCTGTTACCACTCCTACGC | Enh1-3dn-WT | TCATCCAGGGGAATCAGGG | Wt: 335 Mt: 0 |
| | | | Enh1-3dn-MT | GCACAGTGAAAGAGGAGGG | Wt: 0 Mt: 255 |
| ER2 | Enh2-5up | ACCTGAGTCATCAGTACAGTCA | Enh2-3dn-WT | AGTGGAGTGACTGTTCCCT | Wt: 381 Mt: 0 |
| | | | Enh2-3dn-MT | GTTTTTGTGATTTCTTTCCCCCAG | Wt: 0 Mt: 322 (short), 478 (long) |
| ER3 | Enh3-UP-new | AGTTGCAAACCTGAACAGGT | Enh3-3dn-WT | ACCCTGGCTTTTCCAAATCTG | Wt: 346 Mt: 0 |
| | | | Enh3-3dn-MT | TTTGACTGAGGTACAGCAAACC | Wt: 0 Mt: 459 |
| ER2&3 | Enh2-5up | ACCTGAGTCATCAGTACAGTCA | Enh2-3dn-WT | AGTGGAGTGACTGTTCCCT | Wt: 381 Mt: 0 |
| | Enh3-UP-new | AGTTGCAAACCTGAACAGGT | Enh2-3dn-MT | GTTTTTGTGATTTCTTTCCCCCAG | Wt: 0 Mt: 318 |

**Information on Supplementary Datasets**

1. **Suppl. Dataset 1.** Genes with cluster specific gene activities at E14.5 and E17.5 as identified by GeneScore analysis. Cluster identities can be found in Figures 3 and 4.

2. **Suppl. Dataset 2.** E14.5 cluster specific scATAC-seq peaks, associated genes identified by P2G analysis, and DNA motifs for 47 transcription factors present in the scATAC-seq peaks as identified by chromVAR. Column D provides the cluster-specificities of the scATAC-seq peaks, and corresponding identities of these clusters can be found in Figures 3 and 4. Column G provide linked genes identified by P2G link analysis.

3. **Suppl. Dataset 3. E17.5 c**luster specific scATAC-seq peaks, associated genes identified by P2G analysis, and DNA motifs for 47 transcription factors present in the scATAC-seq peaks as identified by chromVAR. Column D provides the cluster-specificities of the scATAC-seq peaks, and corresponding identities of these clusters can be found in Figures 3 and 4. Column G provide linked genes identified by P2G link analysis.

4. **Suppl. Dataset 4.** Differentially accessible peaks between E17.5 and E14.5 by pairwise comparison of corresponding clusters. The clusters compared include nRPCs, tRPCs, PHCs, H&As, and RGCs.

5. **Suppl. Dataset 5**. Differentially expressed genes between E17.5 and E14.5 based on GeneScore by pairwise comparison of corresponding clusters. The clusters compared include nRPCs, tRPCs, PHCs, H&As, and RGCs. Some genes discussed in the text are highlighted.

6. **Suppl. Dataset 6.** Atoh7, Otx2, Pou4f2, and Isl1 CUT&Tag peaks that intersect with E14.5 differential assessable scATAC-seq peaks. Columns A to C are genome coordinates for the CUT&Tag peaks, and columns F to H are genome coordinates of scATAC-seq peaks. Associated Genes identified by P2G are presented in column I. Column J provides cluster specificities of the scATAC-seq peaks. Cluster identities can be found in Figures 3 and 4.

7. **Suppl. Dataset 7.** Enhancers co-bound by Atoh7 and Otx2, and associated genes. Only genome coordinates for Atoh7 CUT&Taq peaks (columns A-C) and the E14.5 scATAC-seq peaks (columns F-H) are presented. Associated genes identified by P2G are presented in column I. Column J provides cluster specificities of the scATAC-seq peaks. Cluster identities can be found in Figures 3 and 4.

8. **Suppl. Dataset 8.** Differential peaks between wild-type and *Atoh7*-null in all corresponding clusters and associated genes as identified by P2G.
